# Single cell analysis of lymphatic endothelial cell fate specification and differentiation during zebrafish development

**DOI:** 10.1101/2022.02.10.479999

**Authors:** Lin Grimm, Elizabeth Mason, Oliver Yu, Stefanie Dudczig, Virginia Panara, Tyrone Chen, Neil I. Bower, Scott Paterson, Kazuhide Okuda, Maria Rondon Galeano, Sakurako Kobayashi, Anne Senabouth, Anne K. Lagendijk, Joseph Powell, Kelly A. Smith, Katarzyna Koltowska, Benjamin M. Hogan

**Author notes:** Authors for Correspondence: Professor Ben Hogan, Organogenesis and Cancer Program, Peter MacCallum Cancer Centre, Melbourne, VIC 3000, Australia, Dr Elizabeth Mason (computational biology lead) Organogenesis and Cancer Program, Peter MacCallum Cancer Centre, Melbourne, VIC 3000, Australia.

## Abstract

During development, the lymphatic vasculature forms as a second, new vascular network derived from blood vessels. The transdifferentiation of embryonic venous endothelial cells (VECs) into lymphatic endothelial cells (LECs) is the first step in this process. Specification, differentiation and maintenance of LEC fate are all driven by the transcription factor Prox1, yet downstream mechanisms remain to be elucidated. We present a single cell transcriptomic atlas of lymphangiogenesis in zebrafish revealing new markers and hallmarks of LEC differentiation over four developmental stages. We further profile single cell transcriptomic and chromatin accessibility changes in zygotic *prox1a* mutants that are undergoing a VEC-LEC fate reversion during differentiation. Using maternal and zygotic *prox1a/prox1b* mutants, we determine the earliest transcriptomic changes directed by Prox1 during LEC specification. This work altogether reveals new transcriptional targets and regulatory regions of the genome downstream of Prox1 in LEC maintenance, as well as showing that Prox1 specifies LEC fate primarily by limiting blood vascular and hematopoietic fate. This extensive single cell resource provides new mechanistic insights into the enigmatic role of Prox1 and the control of LEC differentiation in development.

## Introduction

Lymphatic vasculature plays crucial physiological roles that include the drainage of interstitial fluids, trafficking of immune cells and drainage of dietary lipids. The formation of new lymphatic vessels from pre-existing vessels (lymphangiogenesis) occurs in both development and disease. Signalling through Vegfr3 (Flt4) can be triggered by Vegfc or Vegfd and drives lymphangiogenesis in settings as diverse as development, cancer metastasis, inflammation and ocular disease ^1^. In the embryo, lymphangiogenesis begins when the first LEC progenitors depart the cardinal veins (CVs) from E9.5 in mice and 32 hours post fertilisation (hpf) in zebrafish ^2^. Vegfc-Flt4 signalling drives LEC progenitor sprouting but also up-regulates Prox1 expression in both zebrafish and mice ^3–7^. The transcription factor (TF) Prox1 acts as the master regulator of LEC fate ^8^ and is exclusively expressed in developing LECs in early embryonic vasculature. Loss of Prox1 (Prox1a and Prox1b in zebrafish) leads to a loss of developing lymphatic vessels ^4, 9^. Following departure from the CV, LEC progenitors go on to colonise embryonic tissues and organs and remodel to form functional lymphatic vessels ^10^. While at later stages in mammals there are contributions of LECs from non-venous origins, early embryonic lymphanigogenesis occurs chiefly from the CVs ^11–17^. In mice, both the earliest stages of LEC progenitor sprouting from CVs and maintenance of LEC fate in stable lymphatics are dependent on the function of Prox1 ^9, 18^.

Despite over two decades of study of this enigmatic developmental process, the transcriptomic changes that occur as embryonic venous endothelial cells (VECs) transdifferentiate into LEC progenitors, and further differentiate into mature lymphatics, have not been transcriptionally profiled *in vivo*. In the absence of Prox1 in conditional knockout mice, LECs have been shown to lose the expression of some LEC markers and to gain expression of some blood vascular endothelial cell (BEC) markers^18^. Prox1 is known to autoregulate its own expression and to also regulate Flt4 expression in a positive regulatory loop during early development^7^. Yet how Prox1 controls the transcriptome during LEC specification, differentiation and maintenance has not been described in detail. This is in part because of the technical challenge of accessing early mouse endothelial cells (ECs) in wildtype and mutant embryos, a problem that is not limiting when using the zebrafish embryo.

As recent studies have demonstrated highly conserved expression and function of Prox1 homologues in zebrafish^4, 6, 19, 20^, we here took advantage of the accessibility of the zebrafish embryo to examine developmental lymphangiogenesis using single cell transcriptomics. We provide a resource of new markers of VEC-LEC transdifferentiation, differentiating and mature cell types and validate several new markers with transgenic approaches. We analysed zebrafish zygotic *prox1a* mutants with single cell RNA sequencing and single cell ATAC sequencing. This identified a VEC-LEC fate reversion in the absence of zygotic Prox1a, defined key Prox1-dependent genes in fate maintenance and discovered regulatory regions of chromatin (enhancers) controlled by Prox1. Finally, profiling maternal-zygotic double *prox1a/prox1b* (null) mutant vasculature with single cell RNA sequencing revealed that Prox1 reduces gene expression in early LEC progenitors, including for a core network of conserved haematopoietic and blood vascular fate regulators. Overall, this single cell resource reveals embryonic lymphangiogenesis with Prox1-dependent mechanisms central in specification, differentiation and maintenance of LEC fate.

## Results

### A single cell RNA-seq atlas of embryonic lymphangiogenesis

Zebrafish secondary angiogenesis occurs when Prox1-positive LECs and VECs both sprout from the cardinal vein (CV) in the trunk or the head in a progressive process between ∼32 and ∼48hpf. In the trunk, sprouting LECs migrate dorsally and invest the horizontal myoseptum, where they form a transient pool of parachordal LECs (PLs) from approximately 48hpf^21, 22^. In craniofacial regions of the embryo, LECs sprout from the CVs at several locations^23, 24^. After this, LECs throughout the embryo proliferate and migrate extensively (between ∼56hpf-80hpf) to colonise new regions and tissues^24–27^. In the trunk, LECs anastomose to form the first lymphatic vessels at around 4 days post fertilisation; forming the thoracic duct (TD), dorsal longitudinal lymphatic vessels (DLLVs) and intersegmental lymphatic vessels (ISLVs). In craniofacial regions, they assemble from disparate sources into lateral (LFL), medial (MFL), otolithic lymphatic vessels (OLV) and lymphatic branchial arches (LBA), as well as forming a lymphatic loop (LL) in the head that will later give rise to a unique mural LEC population in the brain (muLECs, also known as FGPs or brain LECs)^28–30^. By 5dpf, major lymphatics in the craniofacial and trunk regions of the embryo are functional and drain dyes and fluids deposited in the peripheral tissues^22, 31^.

To profile stages of development spanning key steps in lymphatic differentiation, we selected: 40hpf, when specification and initial sprouting of LECs are occurring; 3dpf, when immature LECs are migrating through the embryo; 4dpf when LECs are assembling into vessels; and 5dpf when lymphatics are functional and maturing^32^ (**Fig 1a**). We used transgenic zebrafish strains that specifically labelled embryonic vasculature to allow for fluorescence activated cell sorting (FACS, full details in methods). We dissociated whole transgenic embryos and FAC sorted ECs, for profiling on the 10X Chromium scRNA-seq platform. We sequenced 35,634 cells across 3 runs, then merged and normalised the data^33, 34^, filtering low-quality libraries (**Extended data Fig 1a**, full details in methods). To define the cellular identity of each cluster, we systematically evaluated the expression of known markers summarised in **Extended data Table 1a**. We identified 9,771 lymphatic and venous ECs that comprise our single cell atlas of lymphangiogenesis, which is accessible in an interactive CellXGene explorer web app at the link http://115.146.95.206:5006/ (**Fig 1b**, **Extended data Fig 1b-c**).

**Figure 1:**
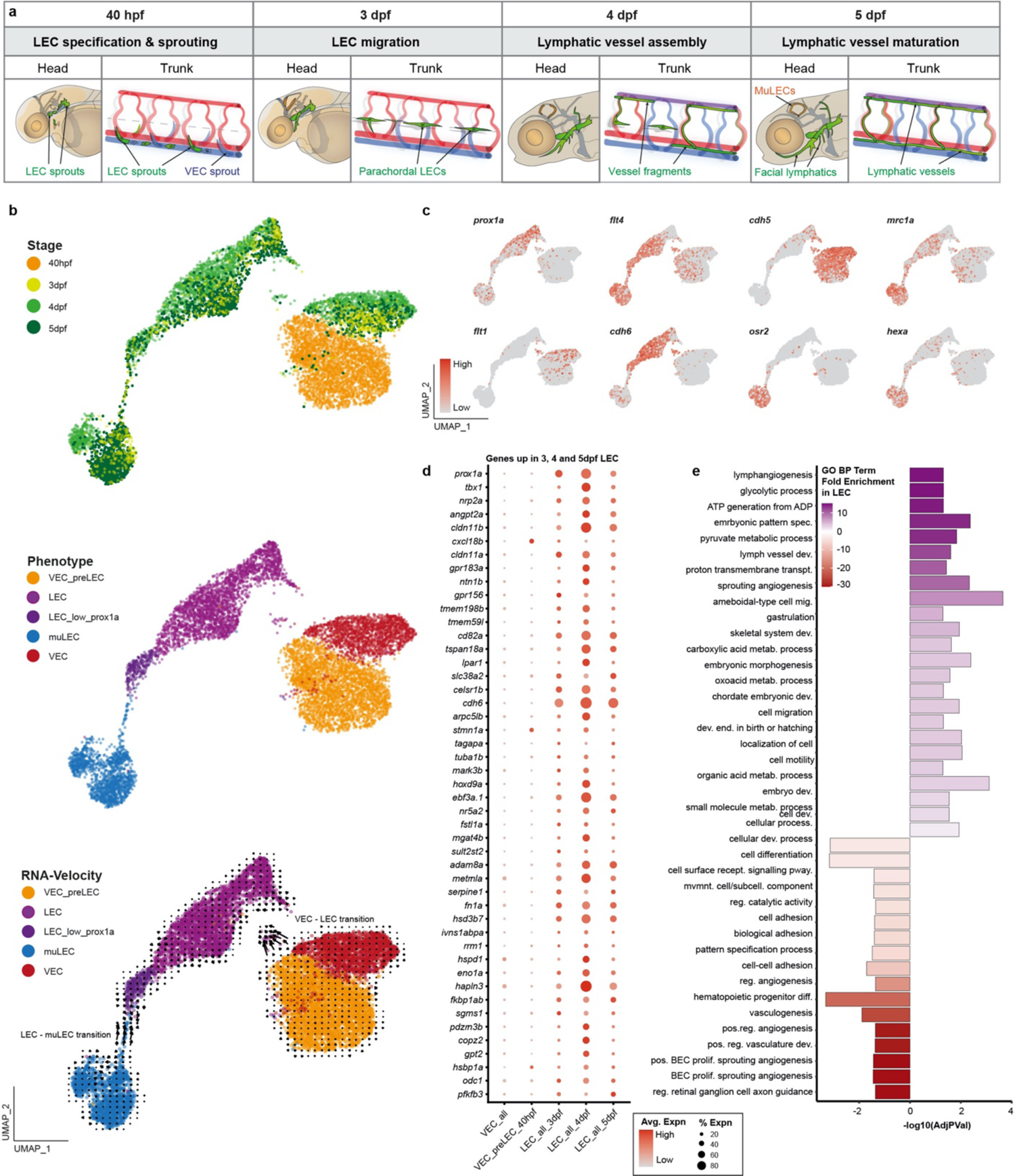
A single cell RNAseq developmental atlas of lymphangiogenesis in zebrafish. **a.** Schematic representation of four key stages of zebrafish lymphatic development in head and trunk: 40hpf encompasses both specification (Prox1-induction) and sprouting; 3dpf migration of LECs; 4dpf assembly of lymphatic vessels; 5dpf maturation functional lymphatics. **b.** UMAP visualization of n=9,771 cells filtered for VEC and LEC populations (n=6 samples; see **Extended data table 1a** for cluster identification and **Extended data Fig 1** for whole dataset) coloured according to developmental stage (top), predicted cell phenotype (middle), and RNA-velocity (bottom). **c.** UMAP visualization of key marker gene expression. Colour scale represents log-normalised expression. LEC markers: *prox1a, cdh6*. BEC markers: *cdh5*. LEC and VEC markers: *mrc1a, flt4*. AEC marker: *flt1*. muLEC markers: *osr2, hexa*. **d.** Dot plot of top genes commonly marking LECs at 3, 4 and 5dpf, with expression displayed in VECs (all stages), VEC_preLEC (40hpf) and LECs at 3, 4 and 5dpf (see also **Extended data table 1b, Extended data Fig 1f**). Colour scale represents average log-normalised expression and point size represents percentage of cells expressing gene. **e.** Bar plot summarizing GO BP analyses of genes DE between 3, 4 and 5dpf LEC and VEC populations (**Extended data table 1c-d**). Y-axis represents enriched BP term, x-axis represents the -log10(*adjusted p value*), bars are coloured and ordered according to fold enrichment of the GO term in LEC. GO terms enriched in LEC are coloured purple, terms enriched in VEC are coloured red. Hpf, Hours post-fertilisation.dpf, Days post fertilisation. BEC, Blood vascular endothelial cell. VEC, Venous endothelial cell. LEC, Lymphatic endothelial cell. muLEC, mural Lymphatic endothelial cell (a.k.a FGP or brain LEC).

This dataset is displayed^35^ as a UMAP in **Fig 1b** coloured according to developmental stage and cell phenotype, with venous (VEC) to lymphatic (LEC) trajectory of differentiation confirmed by RNA-velocity analysis^36^. Comparison of cells at 40hpf with later stages revealed that the earlier populations are transcriptionally distinct from VECs and LECs at 3, 4 and 5dpf. Notably, the 40hpf “VEC preLEC” cluster contained both *prox1a+* and *prox1a-* cells forming a single population rather than discrete clusters. This indicates that LECs are not transcriptionally distinct from VECs at 40hpf, suggesting that while early *prox1a+* cells are specified they have not yet differentiated (**Fig 1c**)^4^. We identified 3 main classes of LECs at 3, 4 and 5dpf: muLECs (1,511 cells) marked by expression of *osr2*^28^, canonical LECs marked by expression of *cdh6* (2,669 cells), and a smaller sub-population of LECs expressing low levels of *prox1a* (LEC_low_prox1a, 310 cells) (**Fig 1c**, **Extended data Fig 1d-e**). To define markers of differentiating canonical LECs at each developmental stages, we applied differential expression (DE) analysis (**Extended data Table 1b**,). This analysis not only captured known LEC markers including *prox1a, angpt2a, tbx1* and *cldn11b*, but also uncovered new genes commonly expressed in canonical LECs across all developmental stages including *grp156, hapln3, cdh6* and *tspan18a* (**Fig 1d, Extended data Fig 1f**). We extended this approach and evaluated global differences between all canonical LECs and all VECs (n=1,240 genes, **Extended data Table 1c**). GO analysis^37^ confirmed the association of biological processes known to be associated with lymphatics with genes upregulated in LEC (n=752 LEC gene set), these terms included “lymphangiogenesis”, “glycolytic process”, “lymph vessel development” and “ameboidal-type cell migration” (**Fig 1e**, **Extended data Table 1d**).

To provide further confidence in the specificity of the new LEC markers defined by this scRNA-seq resource, we used DE analysis to define cluster specific gene expression. We noted that this analysis suggested higher expression of the well-known marker *lyve1b* in muLECs than in LECs, which we validated by examining expression levels of *lyve1b* using the established transgenic line *Tg(lyve1b:DsRed2)* (heat map in **Fig 2a**). We further identified expression of the known kidney epithelial solute transporter *slc7a7* as uniquely expressed in muLEC cells and *fabp11a* as a canonical LEC and VEC marker excluded from muLECs. We generated both new *slc7a7a*–Citrine and a *fabp11a-Citrine* BAC transgenic strains, which confirmed the muLEC and vascular expression patterns and further validated our scSeq atlas (**Fig 2b-c**). Overall, this atlas therefore identifies n=752 specific markers of lymphangiogenesis, the majority of which are new markers (**Extended data Table 1c**), representing new candidate regulators and spanning four developmental stages.

**Figure 2:**
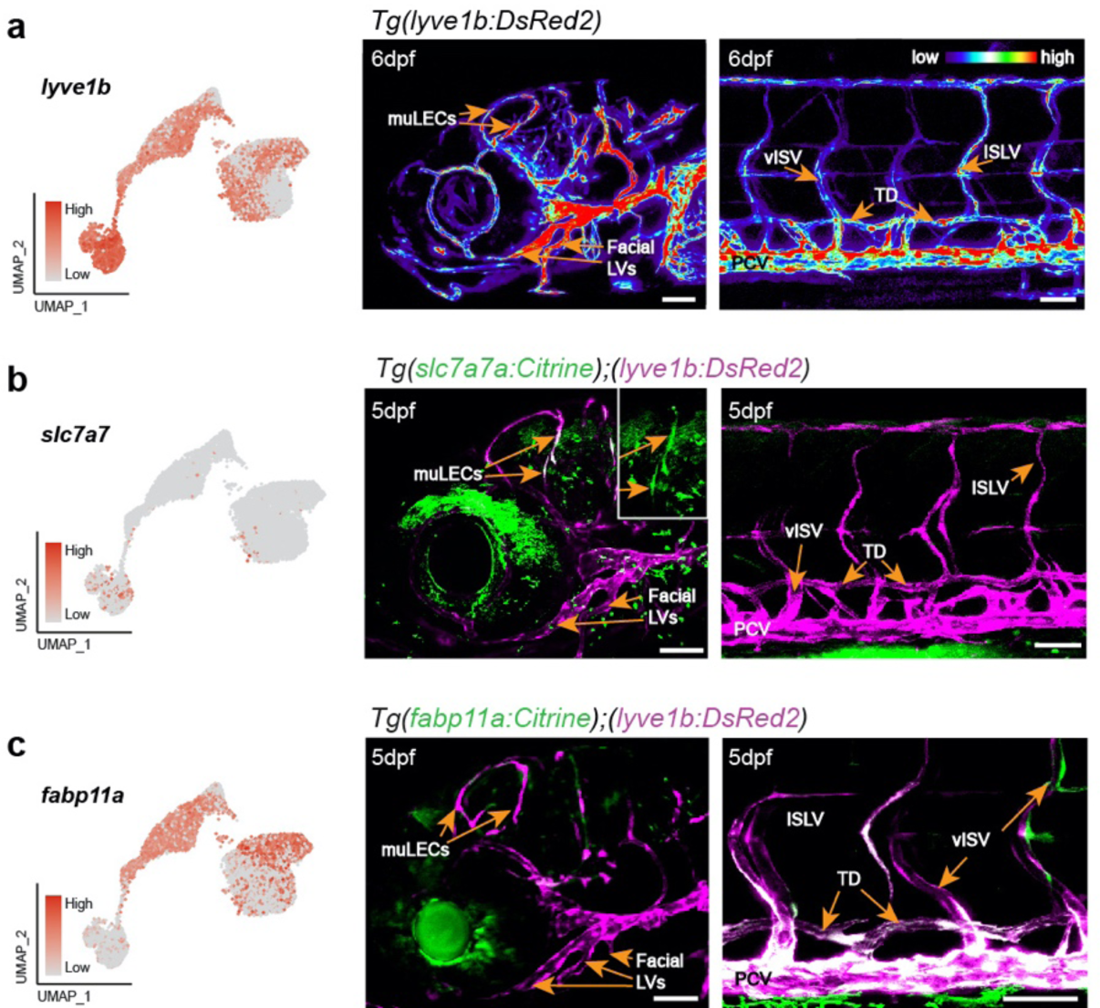
Transgenic marker strains confirm cluster identity and identify vessel specific markers. **a.** UMAP of *lyve1b* expression predicts higher expression in muLECs than LEC populations (left), confirmed by a heat map of a *Tg(lyve1b:DsRed2)* zebrafish larvae at 6dpf showing high *lyve1b* expression in craniofacial (middle) and moderate expression in trunk lymphatic vessels (right). **b.** UMAP of *slc7a7* expression predicts restricted expression in muLEC populations (left), confirmed in lateral confocal projections showing co-expression of *lyve1b* (magenta) and a new BAC transgenic strain for *slc7a7a* (green) in the zebrafish head at 5dpf (middle) which is not expressed in veins and lymphatic vessels of the trunk (right). **c.** UMAP of *fabp11a* expression predicts expression in LEC and VEC but not muLEC populations (left), confirmed in lateral confocal projections showing co-expression of *lyve1b* and a new BAC transgenic strain for *fabp11a* marking venous and lymphatic vessels in the trunk (right) without showing expression in the vasculature of the head (middle) at 5dpf. Lateral confocal images, anterior to the left. muLECs, mural lymphatic endothelial cells; facial LV, facial lymphatic vessels; vISV, venous intersegmental vessel; PCV, posterior cardinal vein; TD, thoracic duct; ISLV, intersomitic lymphatic vessel. Scale bars, 80 µm for head (middle) in (a) and 50 µm for trunk images (right).

### Prox1 maintains LEC identity by repressing blood vascular fate and promoting lymphatic vascular fate at the level of the transcriptome

Prox1 is both necessary and sufficient to drive VEC to LEC trans-differentiation in mammals and this is proposed to occur by Prox1 simultaneously initiating LEC fate and repressing the VEC fate program^8, 18^. Prox1 expression is also necessary for the maintenance of LEC identity during development^18, 38, 39^. Despite the role for Prox1 being well-studied in mouse models, the transcriptomic program controlled by Prox1 *in vivo* has never been profiled. Zebrafish *prox1a* zygotic and maternal zygotic mutants have been previously described^4, 20^. The zygotic mutants retain maternal deposition of *prox1a* in the oocyte, sufficient to drive normal PL formation, LEC migration to the horizontal myoseptum (HM) and initial assembly of lymphatics by 4dpf^4, 19^, however mutants have a reduction in total LEC numbers throughout the face and trunk^4^. When the maternal contribution of *prox1a* is removed, lymphatic development is more severely impaired^4^. We hypothesised that the zygotic mutants which form lymphatic vessels, likely have abnormal vessel identity in the absence of zygotic Prox1. Thus, we applied scRNA-seq to ECs FAC sorted from *Zprox1a^-/-^* mutants^20^ and WT sibling zebrafish at 4dpf, during lymphatic vessel assembly (n=8,075 cells, **Extended data Fig 2a**, **Extended data Table 2a**). Cluster analysis revealed 3 populations of LECs (n=2,068) and a single population of VECs (n=1,051) comprising both mutant and wild type cells, and perhaps most striking, a single population of mutant cells (n=484) transcriptionally similar to VECs, marked by the expression of *aqp1a.1* (**Fig 3a-d**, **Extended data Figs 2b-e**). All three LEC clusters (LEC, LEC_S1, LEC_S2) showed graded expression of lymphatic markers *prox1a*, *cldn11b*, *cdh6* and *angpt2a*, such that “LEC” most closely resembles canonical LECs from our WT atlas. LEC_S1 and S2 represent alternative LEC sub-types with different proliferative potential based on S-phase occupancy and cell cycle marker *mki67* (**Fig 3b,e**, **Extended data Fig 2e**). The expression of *prox1a* was lowest in the more proliferative LEC_S1 cells, and highest in the less proliferative LEC_S2 (**Fig 3b,e**). RNA-velocity analysis suggested a trajectory between the mutant cluster and the LEC cluster (**Fig 3c** lower). Taken together, these findings suggest that the mutant specific cluster sits on a trajectory between LEC and VEC fate, representing cells either failing to fully differentiate from VEC to LEC or LECs undergoing dedifferentiation and fate reversion.

**Figure 3:**
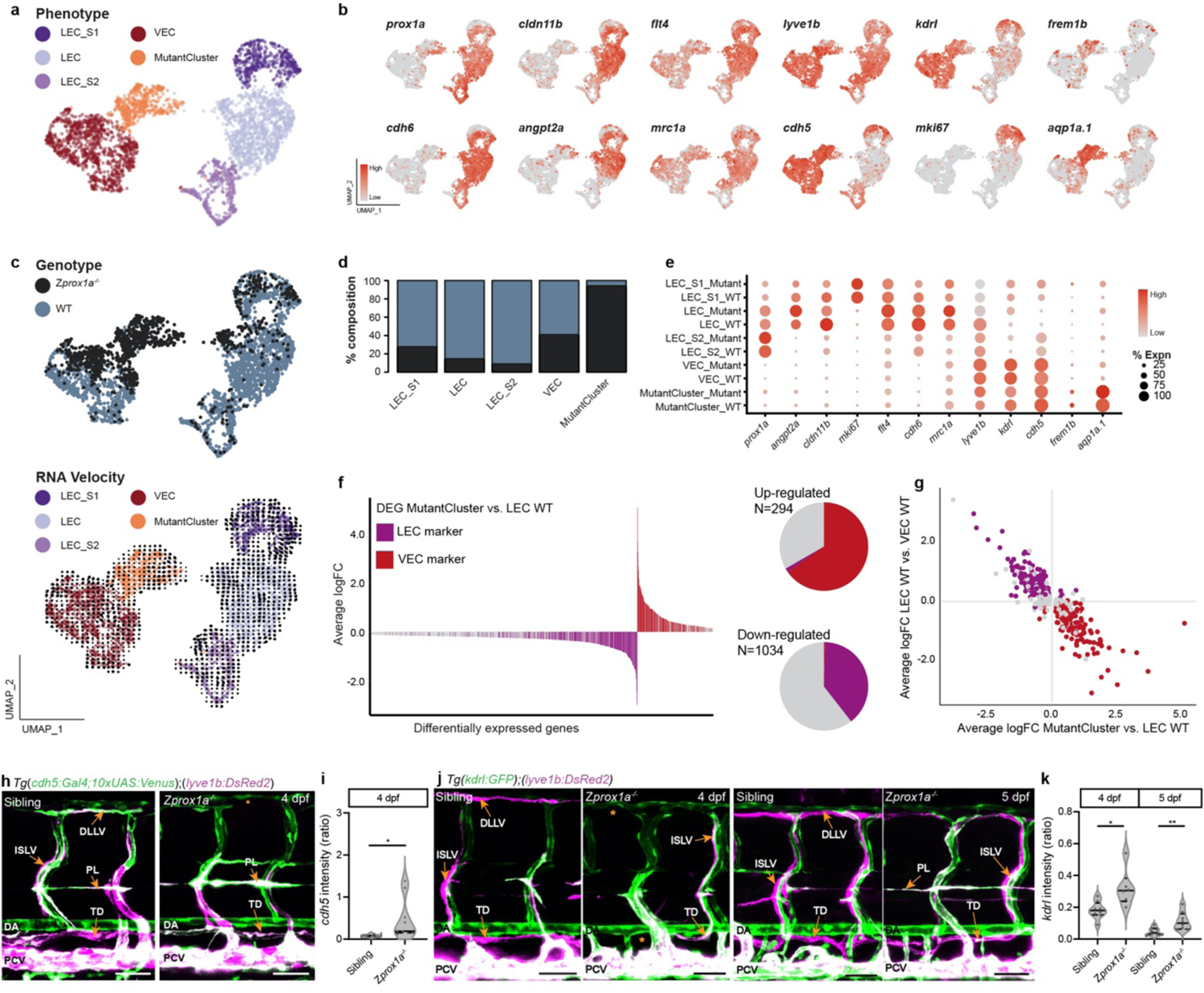
Single cell RNA seq analysis reveals a fate shift from LEC to VEC in the absence of Prox1 in zygotic *prox1a* mutants. **a.** UMAP visualization of n=8,075 cells filtered for VEC and LEC subpopulations (n=2 samples; see **Extended data table 2a** for cluster annotation, **Extended data Fig 3** for full dataset) coloured according to predicted cell phenotype. **b.** UMAP visualization of marker gene expression. Colour scale represents SCT-normalised expression. LEC markers: *prox1a, cldn11b, cdh6, angpt2a*. LEC and VEC markers: *flt4, mrc1a, lyve1b*. BEC markers: *cdh5, kdrl*. Mutant cluster markers: *frem1b, aqp1a.1*. **c.** UMAP visualization with cells coloured according to genotype (*Zprox1a^-/-^* or WT) identifies a mutant specific cluster and RNA velocity analysis suggests a trajectory between the mutant cluster and the LEC cluster, likely indicative of a fate shift. **d.** Stacked bar plot representing the genotype composition of cells in a given phenotypic cluster. **e.** Dot plot of marker expression across defined clusters (indicated on Y-axis). Colour scale represents average SCT-normalised expression and point size represents percentage of cells expressing gene. **f.** Bar plot of average log fold change in gene expression comparing WT LECs with the Mutant Cluster. Y-axis represents average log fold change, x-axis represents differentially expressed genes (Wilcoxin Rank Sum *adjusted p value* < 0.05) and bars are coloured according to status as a LEC (purple) or VEC (red) marker in Fig. 1. Pie charts (right) indicate the LEC and VEC marker composition of genes up-regulated (n=365) and down regulated (n=1,922) in the Mutant Cluster, demonstrating a fate shift. **g.** Concordance in the fate shift between Mutant Cluster and WT LEC with the WT LEC and VEC trajectory. Each point represents significant DE genes (n=2,287) between Mutant Cluster and LEC WT, coloured according to LEC or VEC marker status. X-axis represents average log fold change relative to the Mutant Cluster, Y-axis represents average log fold change relative to LEC WT. **h.** Lateral confocal images of *cdh5* (green) and *lyve1b* (magenta) expression in the developing trunk in WT and *Zprox1a* mutants at 4dpf. **i.** Quantification of *cdh5* intensity in the thoracic duct in WT and mutants (relative to expression in the DA). **j.** Lateral confocal images of *kdrl* (green) and *lyve1b* (magenta) expression at 4dpf (left) and 5dpf (right). **k.** Quantification of *kdrl* intensity in the thoracic duct in WT and mutants (relative to expression in the PCV) at 4 and 5dpf. WT, wildtype. Z, zygotic. TD, thoracic duct, DA, Dorsal Aorta. PCV posterior cardinal vein. ISLV, intersomitic lymphatic vessel. DLLV, Dorsal longitudinal lymphatic vessel. PL, parachordal LEC. Scale bars, 50 µm. *, The error bars represent mean ± s.e.m.; *p=0.01 and **p=0.0094 from an unpaired, two-sided t-test.

To survey how Prox1 maintains normal LEC differentiation, we performed DE analysis comparing the mutant cluster with WT LEC (**Fig 3f**, **Extended data Table 2b**). Overall, a much larger set of genes were downregulated than upregulated in the mutant cluster (n=1,034 vs n=294) with almost half of the most downregulated genes (AvLogFC) highly enriched for LEC markers (eg. *tbx1*, *cdh6*, *cldn11b*). Of the upregulated genes, almost 75% were VEC markers (eg. *sox7*, *kdrl*, *cdh5*) and again this was enriched in the most highly upregulated genes (**Fig 3f, Extended data Fig 2f**). Comparing the change in gene expression between the wildtype LEC cluster and VEC cluster, with the change between wildtype LEC and mutant cluster, we found striking concordance suggesting the mutant cluster is shifted along a LEC fate to VEC fate trajectory (**Fig 3g**). Overall, there is a simultaneous loss of lymphatic fate and re-acquisition of blood vascular gene expression, consistent with work in mouse and with Prox1 function being highly conserved between vertebrates. To validate these observations, we used confocal imaging of Z*prox1a^-/-^* mutants and observed upregulation of blood vascular markers *cdh5* (**Fig 3h-i, Extended data Fig 3a,c**) and *kdrl* (**Fig 3j-k, Extended data Fig 3d-g**) in *lyve1b-*positive lymphatics vessels. We saw a coincident reduction in the expression of *lyve1b* in these vessels (**Extended data Fig 3b).** Interestingly, the difference in *kdrl* levels (relative intensity) in lymphatics between mutants and wildtype was more significant at 5dpf than 4dpf, suggesting that progressive dedifferentiation may be occurring in the mutant (**Fig 3k**). This single cell profile thus demonstrates that Prox1 alone is sufficient to maintain LEC fate and identifies the gene expression maintained by Prox1 during lymphatic differentiation.

### Single cell ATAC sequencing reveals chromatin accessibility signatures in LECs and VECs, identifying lymphatic specific enhancers and predicting key LEC TF families

At the level of gene expression, Prox1 function is essential for VEC to LEC differentiation, however it is unknown if Prox1 controls chromatin accessibility during this process. Thus, we next profiled *Zprox1a^-/-^* mutants at 4dpf during vessel assembly using single nuclei (sn) ATAC-seq (n=3,731 nuclei, **Extended data Fig 4a, Extended data Table 3a**). Cluster analysis (**Fig 4a**) and overall accessibility of key markers (**Fig 4b**) identified similar populations to the scRNA-seq profiling: canonical LEC (LEC_01 n=114 nuclei, LEC_02 n=213 nuclei) and VEC (VEC_01 n=157 nuclei) clusters, and a small but discrete population comprised almost entirely of mutant cells (Mutant Cluster n=47 nuclei). Notably, we found that there was almost no contribution of mutant cells to the LEC clusters (**Fig 4a** lower bar plot), indicating a loss of fate when cell identity is determined at the level of chromatin state **(Extended data Fig 4b)**. Differential accessibility (DA) analysis revealed unique sets of peaks that mark individual VEC, LEC and AEC (arterial EC) clusters (**Fig 4c**).

**Figure 4:**
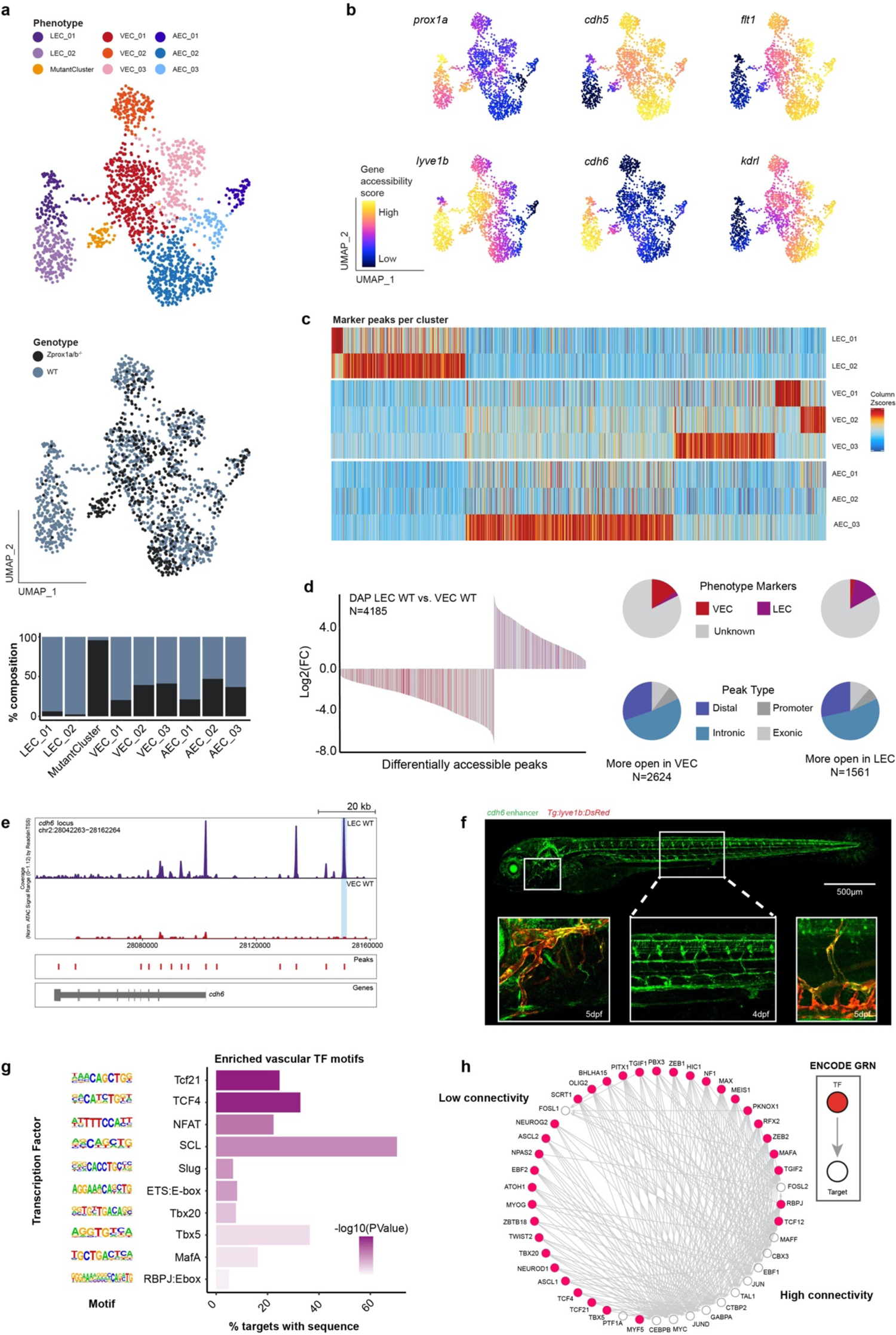
Single nuclei ATAC seq identifies lineage specific regulatory regions in VECs and LECs. **a.** UMAP visualization of snATAC-seq from *Zprox1a^-/-^* mutant and WT endothelial cells at 4dpf (n=3,731 nuclei), coloured by cell phenotype (top) and genotype (middle) (see **Extended data Fig 4** for full dataset). Stacked bar plot (bottom) summarising the composition of wildtype and mutants in each cluster. A mutant specific cluster is identified, similar to Fig 3. **b.** UMAP visualization indicating accessibility of key marker genes (gene accessibility score with imputation) that confirm predicted cluster phenotypes. LEC markers: *prox1a, cdh6*. VEC markers: *cdh5*, *flt1*, *kdrl*. **c.** Heatmap of cluster specific accessible peaks defined using DAP analysis, for all endothelial cells. Colour reflects a column-wise Z-score, rows represent clusters defined in **Extended data Fig 4a** and columns are peaks. **d.** Bar plot of log2 fold change for the differentially accessible peaks (n=4,185 peaks, Wilcoxon Rank Sum, *FDR* < 0.05) between WT LECs and WT VECs. LEC/VEC markers identified in scRNA-seq are coloured in red/purple respectively and demonstrate a strong correlation between chromatin state and the VEC to LEC fate shift. 228/1,561 more accessible peaks in the LEC cluster were associated with LEC markers and 398/2,624 more accessible peaks in VEC cluster associated with VEC markers. Pie charts (right) summarising the proportion of LEC/VEC markers in the differentially accessible peaks (top) and the type of peak (bottom). **e.** Genome accessibility track of LEC marker *cdh6*. Red bars represent peaks in the reproducible peak set from snATAC-seq. The peak highlighted blue indicates a potential enhancer of *cdh6* (*chr2:28150709-28151209*) with significantly more accessible chromatin in WT LECs compared to WT VECs (Wilcoxon Rank Sum, *FDR* < 0.05). **f.** Overall GFP expression of *cdh6* enhancer (*chr2:28150709-28151209*) reporter at 4dpf (top). Lateral confocal image of *cdh6* enhancer reporter in the trunk at 4dpf (bottom middle). Co-expression of *cdh6* enhancer reporter (green) and *lyve1b* (red) in the facial lymphatics (bottom left) and trunk lymphatics (bottom right) at 5dpf. **g.** Vascular TF motifs (n=10 from top n=50) enriched in peaks that are more permissive in WT LEC compared to WT VEC. The depth of bar colour represents the -log10(*RawPVal*), y-axis displays individual motifs and schematic (left) and x-axis represents the percentage of target regions enriched for the motif in the n=1561 peak set more open in WT LEC. **h.** Degree-sorted gene regulatory network displaying known TF binding at genes with more permissive chromatin in WT LECs compared to WT VECs. TFs are represented by red circles (nodes), target genes by white circles (nodes), and known binding of TF to target by a grey arrow (edges). Nodes with a larger number of edges are more highly connected (bottom right), and nodes with fewer edges are less connected (top left). DAP, differentially accessible peaks. FDR, false discovery rate. snATAC-seq, single nuclei ATAC-seq. TF, transcription factor.

To identify phenotype specific chromatin accessibility, we performed DA analysis between WT LEC and WT VEC. This revealed the more accessible regions in the LEC clusters were associated with LEC genes identified in our atlas, and the less accessible regions with VEC genes (**Fig 4d, Extended data table 3b**). DA identified n=1,561 LEC specific peaks and n=2,624 VEC specific peaks representing putative lineage specific enhancers or regulatory elements (**Extended data table 3b**). To test if these regions identified enhancers, we used the zebrafish enhancer detector plasmid system (ZED vector^40^) and tested peaks that were uniquely open 5’ of the newly identified LEC marker gene *cdh6.* To exclude non-specific reporter expression, we also generated F1 embryos using a ZED vector only control. This did display mosaic neuronal and dorsal root ganglion expression **Extended data Fig 4h**, suggesting that the expression from the *cdh6* enhancer in neural structures using this vector should be taken with some caution. No vascular expression was detected. Transgenesis identified one new functional and LEC specific enhancer upstream of *cdh6*, validating the use of this dataset for enhancer discovery (**Fig. 4e-f**).

We next aimed to use this new set of putative enhancer regions in an unbiased manner to identify key TF families likely to regulate LEC development. We performed motif enrichment analysis using HOMER^41^ for all LEC enriched DA peaks identifying n=64 TF family motifs (**Extended data table 3c**). Notably, we found motifs for TCF, ETS, SCL (TAL1), NFAT, TBX, MAF, SLUG and RBPJ family TFs to be enriched in peaks more accessible in LECs (**Fig. 4g**). Analysis of the human homologues in ENCODE data^42^ revealed these TFs regulate a highly connected network of genes expressed in our LEC atlas (**Fig. 4h**). Importantly, a number of these TFs are already known to play important roles in lymphatics (e.g. NFAT^43, 44^, MAFB^45, 46^, TBX1^47^, TCF^48, 49^), supporting the prediction that members of these TF families will play important function roles in LEC development.

### Prox1 reduces the accessibility of chromatin peaks associated with blood vascular and haematopoetic TF motifs

Given that cells in the mutant specific cluster analysed by scRNA sequencing demonstrate loss of LEC gene expression and acquisition of VEC gene expression, we expected chromatin accessibility to change in an equally coordinated manner. However, analysis of accessibility at individual genes (gene score) for the Mutant Cluster revealed changes inconsistent with a simple fate shift (**Extended data tables 4a-b**). Some LEC specific genes (eg. *prox1a*, *cdh6* and *tbx1*) showed loss of transcription in the mutant cluster but increased chromatin accessibility, and some specific VEC genes with increased transcription in the mutant cluster (eg. *cdh5, flt1, gata6*) also showed discordant chromatin changes (**Fig 5a-b, Extended data Fig 4g**). A DAP analysis revealed little concordance between regions with increased accessibility, and expression in LEC or VEC in the scSeq atlas (**Fig 5c**). We identified a subset of genes with more accessible chromatin overall in the Mutant Cluster than either WT LEC or VEC settings (**Fig 5d**). Notably, at the level of individual peaks we found that n=1,726 peaks displayed a striking increase in accessibility in the mutant cluster compared with WT LEC and n=1,794 peaks an increase compared with WT VEC (**Fig 5e**, **Extended data tables 4a-b**). Of these, 431 were common peaks identifying more accessible chromatin regions in the mutant cluster than in either WT LEC or WT VEC (**Fig 5e,** examples in **Fig. 5f**). This suggests that some regions of chromatin open up more than usual during LEC or VEC differentiation in the absence of Prox1.

**Figure 5:**
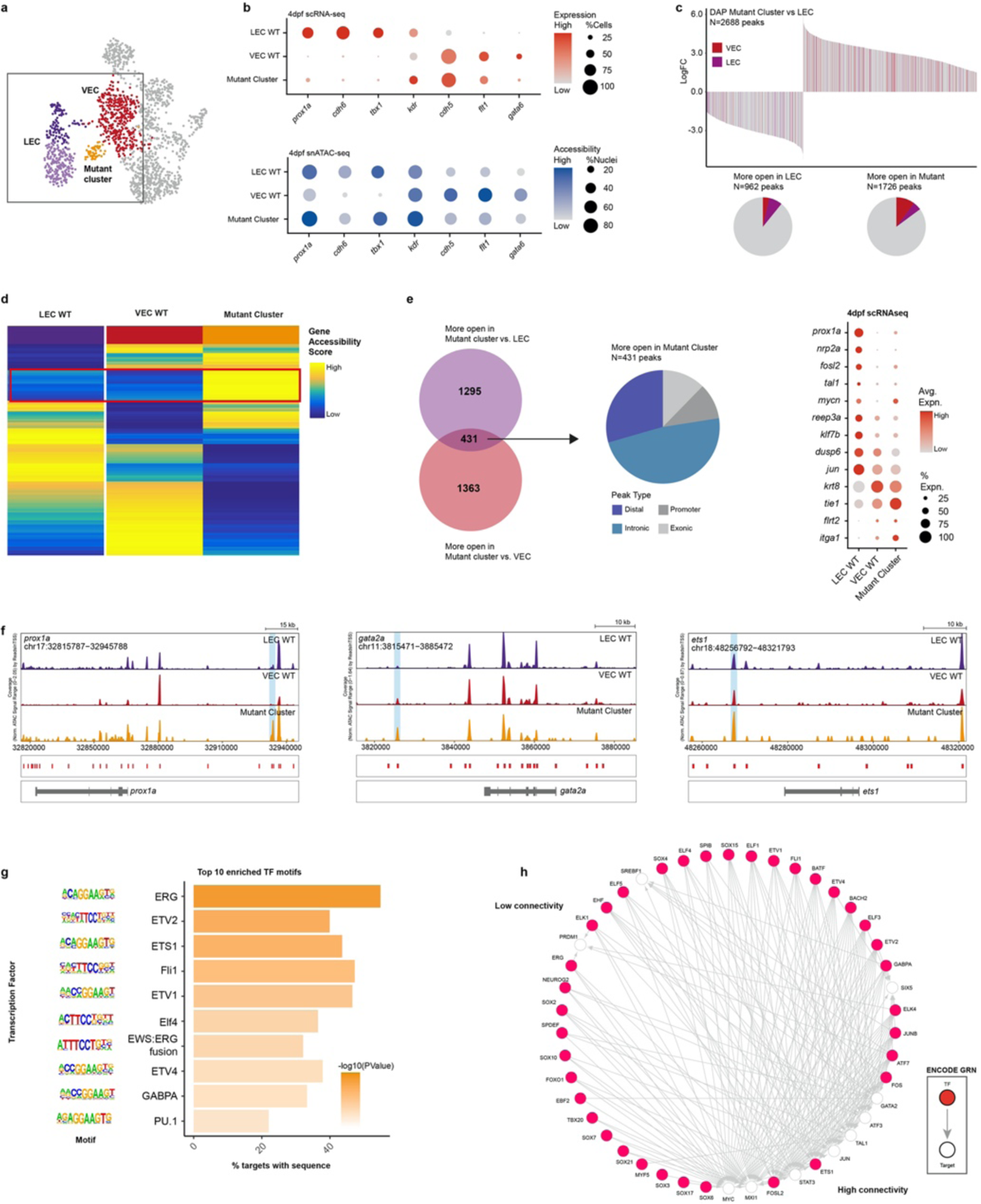
Zygotic *prox1a* mutants display a unique chromatin accessibility state consistent with increased activity of early blood and blood vascular fate transcription factors. **a.** Schematic illustrates DAP analyses between Mutant Cluster vs. LEC WT and Mutant Cluster vs. VEC WT. **b.** Dot plots of scRNA-seq (top) and snATAC-seq accessibility (bottom) data summarising the behaviour of key LEC (*prox1a*, *cdh6*, *tbx1, kdr*) and VEC (*cdh5*, *flt1*, *gata6*) markers in WT LECs, WT VECs and the Mutant Cluster at 4dpf. LEC genes are upregulated (scRNA-seq) and chromatin is more permissive (snATAC-seq) in WT LEC, and VEC genes are upregulated and more permissive in WT VEC. This concordance between gene expression and chromatin accessibility is lost in the Mutant Cluster. The size of the dots represents the proportion of cells or nuclei, and colour represents either SCT-normalised expression or gene score of accessibility. **c.** Bar plot of log2 fold change for the differentially accessible peaks (n=2,688 peaks, Wilcoxon Rank Sum, *FDR* < 0.05) between Mutant Cluster and WT LECs. LEC/VEC markers identified in scRNA-seq are coloured in purple/red respectively and demonstrate regions more open in the Mutant are associated with more vascular than lymphatic genes. Pie charts (bottom) summarising the proportion of LEC/VEC markers in the differentially accessible peaks. **d.** Heatmaps of accessibility (gene accessibility score) for all genes (n=32,020) in WT LECs, WT VECs and Mutant Cluster showing that mutant cluster cells display a unique chromatin state at many genes. Red box indicates genes that are more permissive in the Mutant Cluster than either WT LEC or WT VEC. Colour indicates level of accessibility. **e.** Venn diagram (left) indicates all individual peaks with increased in accessibility in the Mutant Cluster vs LEC or VEC, with n=431 DAP commonly more open than in both LEC WT and VEC WT (*FDR* < 0.05, log2 fold change > 1.5). Pie chart (middle) indicates the proportion of peak types for n=431 DAP classified as distal, intronic, promoter and exonic respectively. Dot plot (right) summarizing the scRNA-seq expression level of n=13 genes with DAP more open in the Mutant Cluster at 4dpf, demonstrating that accessibility changes for these genes did not correlate with changes in transcription. The size of the dot represents the proportion of cells that express the markers in the cluster, and colour represents SCT-normalised expression. **f.** Genome accessibility tracks for key markers with DAP more permissive in the Mutant Cluster: *prox1a, gata2a* and *ets1* (left to right). Red bars represent peaks in the reproducible peak set from snATAC-seq. Blue bars highlight DAP (Wilcoxon Rank Sum, *FDR* < 0.05). **g.** Top 10 enriched motifs (HOMER analysis, *adjusted p value* < 0.05) in the n=431 peaks that are more open in the Mutant Cluster than WT LEC and VEC. The depth of colour represents the -log10(*RawPVal*), y-axis displays individual motifs and schematics (left), and x-axis represents the percentage of peaks enriched for the motif in the n=431 peak set. **h.** Degree-sorted gene regulatory network displaying known TF binding at genes with more permissive chromatin in Mutant Cluster compared to WT LECs. TFs are represented by red circles (nodes), target genes by white circles (nodes), and known binding of TF to target by a grey arrow (edges). Nodes with a larger number of edges are more highly connected (bottom right), and nodes with fewer edges are less connected (top left). DAP, differentially accessible peaks. FDR, false discovery rate. snATAC-seq, single nuclei ATAC-seq. TF, transcription factor.

To investigate the nature of the peaks that were opening in the absence of Prox1, we used an unbiased assessment of TF motifs within these regions. The more open regions were highly enriched for motifs of early acting TFs involved in embryonic vasculogenesis and haematopoiesis, including Erg, Etv2, Etv4, Ets1, Fli1 and Spi1/Pu.1 (**Fig 5g**, **Extended data table 4c**). We examined the human homologues of these TFs in ENCODE^42^ data together with homologues of genes expressed in our atlas and identified a highly connected putative gene regulatory network (GRN) made up of early blood and blood vascular TFs (including ETV2, TAL1, SOX7, SOX17 ERG and other key fate regulators) driving target genes including each other, FOS, JUN, MYC and STAT3 (**Fig. 5h**). We take this to suggest that in the absence of Prox1, TFs that are part of a GRN normally suppressed by Prox1 function are reactivated to drive blood vascular and blood fates. This demonstrates that Prox1 coordinates the correct accessibility of the chromatin and likely controls other TF functions while maintaining LEC fate in ECs. Furthermore, we note that a large number of the genes with increased chromatin accessibility are known regulators of lymphangiogenesis and so this increased accessibility may be a sensitive way to identify key regulatory genes (**Extended data table 4c**).

### Prox1 is required cell autonomously for the normal sprouting of lymphatic progenitors and their contribution to the lymphatic lineage

While the above data identifies the role of Prox1 at stages when it is maintaining LEC identity in assembling LECs, Prox1 is essential for the very earliest decision made when a VEC becomes specified to develop into a LEC^9^. To ask how Prox1 controls the earliest stages of VEC to LEC transdifferentiation, we first needed to generate a complete Prox1 loss of function model in zebrafish. We generated *prox1a^+/-^*, *prox1b^+/-^* double heterozygous animals (lacking both Prox1 homologues) and then used germline transplantation approaches to produce animals carrying a double mutant germline (**Extended data Fig 5a**). Genetic crossing of these animals generated embryos that were maternal zygotic (MZ) *MZprox1a^-/-^*, *MZprox1b^-/-^* mutant zebrafish completely lacking *prox1a/b* transcript expression or maternal deposition (**Fig 6a; Extended data Fig 5b-c**). A quantitative phenotypic analysis revealed double MZ mutants show a severe reduction of facial lymphatics and a near complete loss of lymphatic vessels in the trunk by 4dpf (**Fig 6e**, **Extended data Fig 5d-g**) ^4^. Furthermore, *MZprox1a^-/-^*, *MZprox1b^-/-^* mutants initially show a delay in the formation of PLs as these cells emigrate the CV to invest the HM, but despite this delay, these cells eventually seed the HM before failing to undertake any further migration (**Fig 6d,f**). We saw evidence for genetic interaction between *prox1a* and *prox1b* in PL formation and no change in the overall number of cells sprouting from the PCV or number of cells contributing to the venous ISVs (blood vessels) suggesting both *prox1a* and *prox1b* contribute specifically to early PL development (**Fig 6g-k)**. To test whether LECs require Prox1a in a cell autonomous manner, we performed embryonic transplantation to generate chimeric embryos. Due to challenges generating and maintaining large numbers of double MZ mutant embryos, we performed transplantion of *MZprox1a^-/-^* mutant cells into wildtype hosts only and assessed the contributions of vascular grafts to arteries, veins and lymphatics (**Extended data Fig 6a**). MZ*prox1a^-/-^* mutant cells efficiently contributed AECs, VECs but not LECs to developing vessels in otherwise wildtype hosts at 5dpf (**Extended data Fig 6b-d**). Thus, we confirmed that Prox1 is necessary cell autonomously for zebrafish lymphatic development and that the *MZprox1a^-/-^*, *MZprox1b^-/-^* mutant phenotype is more severe than any previously described zebrafish Prox1 mutants.

**Figure 6:**
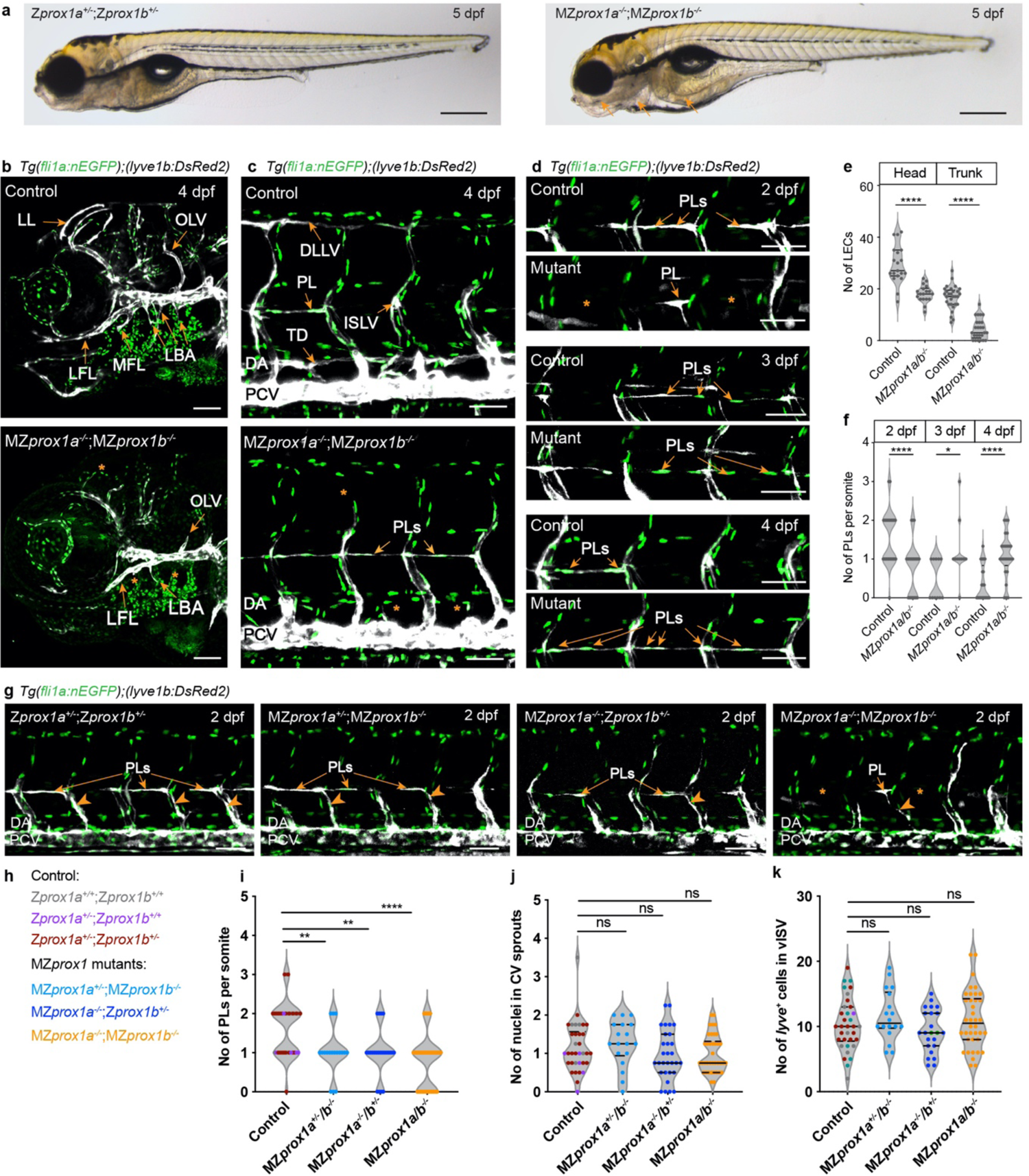
maternal zygotic *prox1a;prox1b* double mutants display a specific loss of lymphatic vessels throughout the developing embryo. **a.** Overall morphology of control and *MZprox1a^-/-^;MZprox1b^-/-^* mutants at 5dpf. Arrows indicate oedema around eyes, heart and intestine. Scale bars 500 µm. **b.** Lateral confocal images of zebrafish heads at 4dpf showing endothelial cell nuclei (green) and venous and lymphatic vessels (white) in control (upper) and *MZprox1a^-/-^; MZprox1b^-/-^* embryos (lower). Facial lymphatics are absent or shorter or missing (asterisk) in the mutants (asterisk). Scale bar, 80 µm. **c.** Lateral confocal images of zebrafish trunks at 4dpf in control (upper) and *MZprox1a^-/-^; MZprox1b^-/-^* mutants (lower), showing absent lymphatic vessels (asterisk) but retained PLs (arrows in lower). Scale bars, 50 µm. **d.** Lateral confocal images of PLs in the horizontal myoseptum at 2, 3 and 4dpf. PLs form later in *MZprox1a^-/-^;MZprox1b^-/-^* mutants and accumulate in the horizontal myoseptum while the PLs of control embryos emigrate the HM by 4dpf. Scale bars, 50 µm. **e.** Quantification of lymphatic endothelial cell (LEC) number from 4dpf heads control: n=19, *MZprox1a^-/-^;MZprox1b^-/-^* mutants: n=18) and trunks (control: n=29, *MZprox1a^-/-^; MZprox1b^-/-^* mutants:n=25). Error bars represent mean ± s.e.m.; **** p < 0.0001, from an unpaired, two-sided t-test. **f.** Quantification of number of PLs per somite in trunks at 2dpf in controls (n=41) and *MZprox1a^-/-^;MZprox1b^-/-^* mutants (n=38), at 3dpf in controls (n=11) and *MZprox1a^-/-^; MZprox1b^-/-^* mutants (n=10) and at 4dpf in controls (n=29) and *MZprox1a^-/-^; MZprox1b^-/-^* mutants (n=25). Error bars represent mean ± s.e.m.; **** p < 0.0001, * p < 0.05, from an unpaired, two-sided t-test. **g.** Lateral confocal images showing defects in the formation of PLs in mutants upon loss of *prox1a* compared with controls in the trunk. Genotypes are indicated. **h.** Colour coded list of analysed genotypes abbreviated in (**i-k**). Each embryo has a defined genotype for *prox1b* which is represented in colour code as displayed in j. **i.** The number of PLs formed (per somite) at 2dpf in genotypes indicated. Decreasing gene dosage for *prox1a* and *prox1b* progressively reduced the initial seeding of the HM by PLs. **j.** The number of cells in sprouts departing the PCV is unchanged in mutants showing that sprouting occurs normally. **k.** The number of cells in vISVs is unchanged in mutants indicating no effect on the venous endothelium. LL, lymphatic loop. OLV, otholitic lymphatic vessel. LFL, lateral facial lymphatic vessel. MFL, medial facial lymphatic vessel. LBA, lymphatic branchial arches. DLLV, dorsal longitudinal lymphatic vessel. PL, parachordal LEC. ISLV, intersomitic lymphatic vessel. TD, thoracic duct. DA, dorsal aorta. PCV, posterior cardinal vein.

### Prox1 functions to suppress blood and blood vascular fate during the early specification of LEC fate in the embryo

To understand the very earliest role that Prox1 plays in LEC development, we profiled the endothelium of *MZprox1a^-/-^*, *MZprox1b^-/-^* mutant zebrafish at 40hpf (when cells are both being specified to LEC fate and also actively sprouting from various regions of the PCV^3, 4, 6^) using single cell RNA seq as described above (**Extended data table 5a, Extended data Fig 7a-c**). Using key marker expression and DE analysis, we identified populations of cells that included VECs, endocardium, mixed populations of VECs and Prox1+ LEC progenitors (named LEC_VEC) and clusters with expression of some VEC and AEC markers that were likely still differentiating (**Fig 7a-c, Extended data Fig 7d-h**). Based on expression of known markers and genes associated with EC sprouting (*mki67*, *pcna*), we defined two populations of secondary sprouts of venous origin (LEC VEC 01 n=713 cells, LEC VEC 02 n=677 cells) and a single population of cells representing the cardinal vein (PCV n=812 cells; **Fig 7d-e**). Consistent with our observations in **Fig 1,** cells expressing *prox1a* at 40hpf were not transcriptionally distinct from sprouting VECs and failed to form a “lymphatic progenitor” cluster, suggesting they are specified and express *prox1a* but not yet differentiated. RNA velocity analysis suggested that these three clusters remained closely related, consistent with little differentiation between these populations at this stage (**Fig 7a** right).

**Figure 7:**
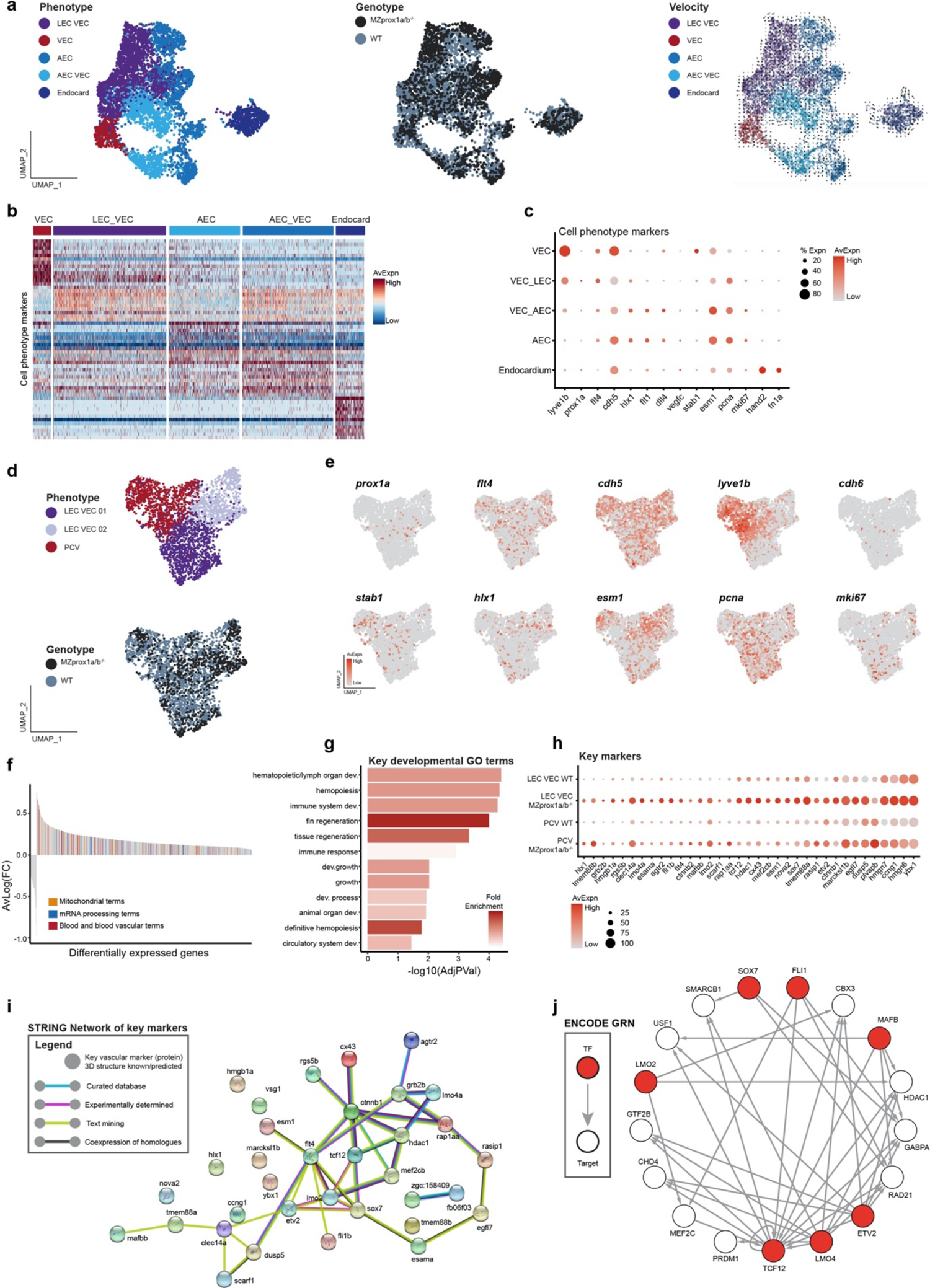
Single cell profiling of maternal zygotic *prox1a;prox1b* double mutants reveals that Prox1 homologues initially suppress gene expression to drive a fate transition. **a.** UMAP visualisation of endocardial, venous and arterial endothelial cells (Level 02 n=5,347) coloured according to predicted cell phenotype (left), genotype (middle) and RNA-velocity (right). **b.** Heatmap displaying expression of phenotype specific genes defined using differential expression analysis. Columns are cells grouped by phenotype assignment, rows are genes, and colour indicates the average level of expression. **c.** Dot plot indicating expression of key markers used to define cell phenotype. Colour scale represents average SCT-normalised expression and point size represents percentage of cells expressing gene. **d.** UMAP visualisation of venous endothelial cells (Level 03 n=2,747) coloured according to predicted cell phenotype (top) and genotype (bottom). **e.** UMAP visualisation of key marker gene expression. Colour scale represents SCT-normalised expression. **f.** Bar plot indicating average log fold change of n=1,186 significantly different genes between all *MZprox1a/b^-/-^* and WT venous endothelial cells (LEC VEC 01, LEC VEC 02 and PCV combined) at 40hpf (Wilcoxin Rank Sum *adjusted p value* < 0.05). Colour indicates genes associated with one or more mitochondrial, mRNA processing and blood and blood vascular GO terms. **g.** Bar plot summarizing GO BP analysis of the n=1,137 genes upregulated in the *MZprox1a/b^-/-^* venous endothelial cells. Y-axis represents enriched GO BP term, x-axis represents the -log10(*adjusted p value*) and bars are coloured according to fold enrichment. **h.** Dot plot of n=36 key blood vascular markers upregulated in the *MZprox1a/b^-/-^* venous endothelial cells, indicating genotype specific expression in LEC VEC and PCV cell phenotypes. Colour scale represents average SCT-normalised expression and point size represents percentage of cells expressing gene. **i.** STRING analysis of n=36 key blood vascular markers upregulated in the *MZprox1a/b^-/-^* venous endothelial cells. **j.** Degree-sorted gene regulatory network displaying known TF binding at n=1,137 genes upregulated in the *MZprox1a/b^-/-^* venous endothelial cells. TFs are represented by red circles (nodes), target genes by white circles (nodes), and known binding of TF to target by a grey arrow (edges). Nodes with a larger number of edges are more highly connected (bottom right), and nodes with fewer edges are less connected (top left).

To identify the earliest transcriptional changes controlled by Prox1 in lymphatic development, we used DE to evaluate the global differences between *MZprox1a^-/-^*, *MZprox1b^-/-^* and WT in the LEC VEC 01, LEC VEC 02 and PCV clusters of sprouting cells (**Fig 7f, Extended data table 5b**). We found a significant upregulation of a large set of genes in the absence of Prox1 that were identified as enriched for blood vascular and haematopoietic genes by GO-term and marker analysis (**Fig 6g-h,** n=1,137 genes, **Extended data table 5c**). We also saw increased expression of mitochondrial metabolism genes and of pre-mRNA splicing genes in the absence of Prox1 (**Fig. 7f, Extended data table 5c**). At this stage of development, in contrast to the 4 dpf stage, we saw little evidence of positively regulated genes downstream of Prox1 (just n=49 genes downregulated in *MZprox1a^-/-^*, *MZprox1b^-/-^*). We next examined the genes up-regulated in Prox1 mutants that we identified as key early blood and vascular developmental regulators. This includes *esm1, flt4, mef2c, hdac1, lmo2, lmo4, rasip1, egfl7, dusp5, clec14a, fli1b* and others (**Fig. 7h-i**). STRING^50^ analysis suggested the presence of inter-related gene networks and so we further examined up-regulated TFs in this network by leveraging human homologues in ENCODE data and identifying known target genes expressed in our dataset. This identified a GRN containing SOX7, FLI1, LMO2, ETV2, TCF12, LMO4 and MAFB as well as known targets of these TFs, that is repressed in response to Prox1-function during LEC specification (**Fig. 7j**). This is highly concordant with observations at later stages of up-regulated TF activity in 4-day old mutants (**Fig. 5**). Overall, we believe this suggests that in the earliest stages of LEC specification Prox1 is not actively driving a fate program but rather is acting primarily to down-regulate blood vascular and hematopoietic fate. It seems likely that the negative regulation of core blood and blood vascular fate genes (those in **Fig. 7j**), would be sufficient to drive the differentiation in these earliest specified LECs.

## Discussion

In this study, we first present a single cell RNA seq analysis of four key stages of embryonic lymphangiogenesis in a vertebrate embryo, revealing new markers and potential regulators of LEC differentiation. We further profile the ECs of *Zprox1a* mutants, which form lymphatic structures, but these “lymphatics” dedifferentiate or revert their fate to blood vascular in a mutant specific, fate-shifted, single cell cluster. This analysis identified the transcriptional code maintained by Prox1 in order to maintain LEC identity and also shows the highly conserved nature of Prox1 function comparing zebrafish with mice. Overall, the striking fate shift identified across the whole transcriptome confirms that the function of just one transcription factor (Prox1) is sufficient to maintain cellular identity, validating its status as the master regulator of LEC identity.

As well as single cell profiling of the developing LEC transcriptome, we also performed snATAC seq on 4 dpf zygotic *prox1a* mutants and wildtype, again identifying a mutant specific, fate shifted cluster in this analysis. Analysis of the wildtype LECs and VECs revealed strong concordance between chromatin accessibility at LEC enhancers and the transcriptional profile for LEC and VEC specific genes. Interestingly however, analysis of the mutant cluster chromatin identified ectopically open regions and peaks with a distinct discordance between chromatin accessibility at enhancers and transcriptional profile. In the mutant cluster at 4dpf, a large number of opened chromatin regions were enriched for TF motifs for earlier acting vasculogenic and haematopoietic TFs. This suggests that regulators of earlier vascular fate become more active and increase chromatin accessibility at specific targets in the absence of Prox1. Overall, this revealed that at the level of chromatin accessibility the fate-shifted ECs display a more immature state, perhaps a consequence of regulatory “confusion” due to a failed fate transition. It seems likely that Prox1 has a combinatorial function as part of a larger GRN of developmental TFs, which has yet to be studied in detail.

Additional biological insights from this study come from our analysis of double maternal zygotic *prox1a, prox1b* mutants. These mutants are presumed “null” mutants for Prox1 orthologues and they revealed (through scRNA seq) that during its earliest role in vascular development, in LEC fate specification and VEC-LEC transdifferentiation, Prox1 functions primarily to negatively regulate blood and blood vascular fate. While this may represent a downstream program rather than direct repression of gene expression by Prox1 orthologues, it is notable that Prox1 and *Drosophila* Prospero have been reported to be able to function as repressors in a context dependent manner ^51–55^. Interestingly, the analysis of upregulated genes identified a set of TFs that are known early regulators of blood and blood vascular fates during embryonic haematopoiesis and vasculogenesis. These included Sox7, Etv2, Lmo2 and Lmo4 and we take this to suggest that Prox1 functions during VEC-LEC transdifferentiation at least in part by blocking expression of early acting blood vascular fate driving TFs. This is in line with maintaining a negative regulatory relationship with these and other fate driving TFs at 4 dpf as described above. It will be interesting in the future to understand at a mechanistic level if Prox1 is actively repressing gene expression at bound targets to block alternative cell fates and if so, if the activity of Prox1 switches between repressor or activator at different stages or target genes.

Altogether this study describes the process of developmental venous to lymphatic transdifferentiation, early LEC differentiation and LEC maintenance in detail, *in vivo*. We describe the role of Prox1 during this process, revealing a conserved and dynamic regulatory process with unprecedented resolution. This resource will help to understand lymphangiogenesis in contexts beyond the embryo, such as in pathological lymphangiogenesis in metastasis, inflammation and tissue repair. Finally, we suggest that combining zebrafish mutant models, with their inherent ease of cellular accessibility, and the resolution afforded by single cell profiling, will be an approach that yields powerful new insights in many areas of developmental biology in the future.

## Acknowledgements

This project was supported in part by NHMRC Ideas grant 2004300 and ARC Discovery project DP180102846. B.M.H. was supported by NHMRC Fellowship 1155221. We thank Angelika Christ and the IMB Sequence Facility (University of Queensland), Tim Semple and Peter Mac Genomics Facility, the Research Computing Facility at Peter Mac, FACS facilities at UQ and Petermac and Isaac Virshup for assistance.

## Author contributions

L.G. and E.M. performed, analysed experiments and co-wrote manuscript. B.M.H, K.K and N.L.H conceptualised experiments, analysed data and co-wrote manuscript. S.D, T.C, O.Y, N.I.B, S.P, K.O, A.S and A.L. performed and analysed experiments. J.P, and K.A.S, provided key unpublished reagents and analysed data.

## Conflicts of interest

The authors declare no conflict of interests.

### Online Methods Zebrafish husbandry

Zebrafish work was conducted in compliance with animal ethics committees at the Peter MacCallum Cancer Centre, the University of Melbourne and the University of Queensland. Published transgenic lines used were: *Tg(fli1a:nEGFP)^y7^* ^56^*; Tg(−5.2lyve1b:DsRed)^nz1^*^01^ ^24^; and *Tg(kdrl:Has.HRAS-mCherry)^s9^*^16^ ^21^. Published mutant lines used were *prox1a^i278^* ^20^, *prox1b* ^4, 19^. *Tg(lyve1b:Venus)^uq51bh^* was generated as previously described^28^ but here using an independent genomic integration with the same construct.

### Generation of maternal zygotic mutants

Germline replacement was performed using embryonic transplantation as previously described ^4, 57^. Maternal zygotic (MZ) *prox1a* mutant embryos were made by crossing germline replaced *prox1a^i278−/−^*;*prox1b^sa0035+/−^* females with *prox1a^i278+/−^*;*prox1b^sa0035+/−^* males ^4, 57^. Genotyping of individual embryos during transplantation and phenotypic analysis was performed as previously described ^4^.

### Transgenesis, genome-editing and genotyping

All microinjections were performed as previously described ^58^. *slc7a7a^BAC^:slc7a7a-Citrine^uom10^* and *fabp11a^BAC^:fabp11a-Citrine^uom10^* recombineering was performed as previously described^59^.

Primers for BAC recombineering:

*slc7a7a*-BAC-Citrine-forward:5’-AACTGCTTTAGACAGTGTTTTTTGGTACCATCCCATATATTTAAAAAACAGCCACCATGGTGAGCAAGGGCGAGGAG-3’

*slc7a7a*-BAC-Citrine-reverse:5’-TTCGACACCTCAGGGGATGCCTCTTCTGCAGGCGTAGGGCTGTAGGACGCTCAGAAGAACTCGTCAAGAAGGCG-3’

*fabp11a*-BAC-Citrine-forward:5’-TTACAGCTGTTGCGAGATTGAAAAGTAGAGGAGCATCATTATTCGGGAAAGCCACCATGGTGAGCAAGGGCGAGGAG-3’

*fabp11a*-BAC-Citrine-reverse:5’-TCAAAGTTGTCGCTGGTGGTCATTTTCCACGTTCCTACGAATTTGTCAACTCAGAAGAACTCGTCAAGAAGGCG-3’

### Enhancer detection transgenesis and analysis

A 501bp PCR fragment of *cdh6* enhancer (*chr2:28150709-28151209*) was cloned into the zebrafish enhancer detection (ZED) vector^40^. Empty ZED vector was injected as previously described^40^. Briefly, 1nL of construct at 40 ng/μl or 45ng/μL and tol2 transposase mRNA at 100 ng/μl or 55ng/μl was injected into the one-cell stage wild type zebrafish embryos. All F0 embryos were screened for skeletal muscle DsRed2 expression. Stable F1 embryos were imaged on a Zeiss LSM 780 or Leica TCS SP8 DLS microscope. Images were processed using ImageJ 2.0.0. or 2.3.0.

### Imaging and quantification

Imaging was conducted at the Centre for Advanced Histology and Microscopy (Peter MacCallum Cancer Centre). Imaging of live samples was performed using a Zeiss LSM 710 FCS confocal microscope, a Zeiss LSM 780 FCS confocal microscope, or an Olympus FV3000 confocal microscope. Mounting and imaging were performed as previously described ^60^. In **Figs 6e-f,i-k**, and **Extended data Fig 5f-h, k-m** quantification of vascular phenotypes was performed as previously described ^60, 61^. In Figs 3h-k, *cdh5:Venus* or *kdrl:GFP* intensity in the TD was measured using Imaris software (Bitplane) and normalised to fluorescence intensity of the DA for *cdh5* and to the fluorescence intensity PCV for *kdrl* in the same embryos.

### Fluorescence activated cell sorting

FACS was performed at the Peter MacCallum Cancer Centre and the University of Queensland. For **Fig 1**, we isolated cells using the following transgenic lines: 40hpf, *Tg(fli1a:nEGFP)^y7^*; 3,4 and 5dpf, *Tg(−5.2lyve1b:Venus)^uq47bh^, Tg(kdrl:Has.HRAS-mCherry)^s9^*^16^ For **Fig 3** and **Fig 4**, we isolated cells using the following transgenic lines: 4dpf, *Tg(fli1a:nEGFP)^y7^, Tg(−5.2lyve1b:DsRed)^nz101^.* For **Fig 6**, we isolated cells using the following transgenic lines: 40hpf, *Tg(fli1a:nEGFP)^y7^.* To dissociate embryos and obtain single cell suspensions, we followed published protocols ^62^. Briefly, at the desired developmental stage we deyolked embryos by pipetting up and down and rinsing in calcium free ringers solution, we centrifuged at 2000rpm for 5’ at 4°C and remove supernatant and dissociated the cells by incubating in liberase [2.5mg/mL] (Cat #5401119001 Sigma-Aldrich) diluted at a 1:35 ratio in DPBS at 28.5°C for approximately 5’, homogenizing the samples during and after the incubation. To stop the reaction, we added CaCl2 to a final concentration of 1-2mM and FBS to a final concentration of 5-10%. We centrifuged at 2000rpm for 5’ at 4°C. discarded the supernatant, in order to be able to asses live vs. dead cells, we re-suspend the cells solution in Zombie Violet TM Viability Dye (Cat# 423113, BioLegend) and incubated for 20’ at RT softly rocking, we rinsed the cells by centrifuging and resuspending in DPBS/EDTA, for ATAC-seq experiments samples were resuspended in 2%BSA/PBS. Suspension was filtered through a strainer and taken to the FAC sorting facility. In the Flow Cytometry facility, we used the BD FACS Aria Fusion sorter (BD Biosciences), we based the selection for the desired population on FSC and SCC, alive cells were selected based on the Zombie Violet profile and double positive cells for the desired transgenics were targeted according to the expression profiles of single cells. Double positive cells were sorted in 300uL 100% FBS in a cold block and taken immediately to the sequencing facility.

### scRNA-seq library preparation

Library preparation and sequencing was performed at the Institute for Molecular Bioscience Sequencing Facility (University of Queensland) or Peter Mac Genomics Facility. Single cell suspensions were sorted by FACS, spun down to concentrate and a cell count was performed to determine post-sort viability and cell concentration. Single cell suspension was partitioned and barcoded using the 10X Genomics Chromium Controller (10X Genomics) and the Single Cell 3’ Library and Gel Bead Kit (V2 10X Genomics PN-120237; V3.1; 10X Genomics; PN-1000123). The cells were loaded onto the Chromium Single Cell Chip A (10X Genomics; PN-120236), B (10X Genomics; PN-1000073 or PN-1000074) or G (10X Genomics; PN-1000120) to target 10,000 cells. GEM generation and barcoding, cDNA amplification, and library construction was performed according to the 10X Genomics Chromium User Guide. The resulting single cell transcriptome libraries contained unique sample indices for each sample. The libraries were quantified on the Agilent BioAnalyzer 2100 using the High Sensitivity DNA Kit (Agilent, 5067-4626). Libraries were pooled in equimolar ratios, and the pool was quantified by qPCR using the KAPA Library Quantification Kit - Illumina/Universal (KAPA Biosystems, KK4824) in combination with the Life Technologies Viia 7 real time PCR instrument. After the initial sequencing run, libraries were re-pooled according to estimated captured cells as determined using the Cell Ranger software (10X Genomics).

### Sequencing of scRNA-seq libraries

At the IMB(UQ) genomics facility, denatured libraries were loaded onto an Illumina NextSeq-500 and sequenced using a 150-cycle High-Output Kit as follows: 26bp (Read1), 8bp (i7 index), 98bp (Read2). Read1 supplies the cell barcode and UMI, i7 the sample index, and Read2 the 3’ sequence of the transcript. At the Peter Mac Molecular Genomics facility, single cell transcriptome libraries were sequenced on an Illumina NovaSeq 6000 using S4 300-cycle chemistry. Read1 supplies the cell barcode and UMI, i7 the sample index, and Read2 the 3’ sequence of the transcript. Sequencing read lengths were trimmed to 28bp (Read1), 8bp (i7 index), 91bp (Read2), ensuring compatibility with the 10X Genomics analysis software, Cell Ranger.

### scRNA-seq data processing and analysis

All code and documentation associated with this analysis is publicly available under an open source software license at: https://atlassian.petermac.org.au/bitbucket/users/tyrone.chen/repos/hogan_lab/browse/2022_Grimm_Mason_et_al_PROX1/

Relevant functions are in italics for reference. Where necessary fastq files were made using *Cell Ranger* ^63^ (version 3.1.0 or 3.0.2) *mkfastq*. Sequencing QC was assessed using *FastQC* 0.11.6 and *MultiQC* viewer for aggregated reports. *Cell Ranger count* and *aggr* were used to generate aggregated count files mapped to GRCz11 (Ensembl 101), without depth normalisation. Doublets were identified from the filtered aggregated count files using *Scrublet* ^64^ in *Python* version 3.6 and filtered from subsequent analyses. For the *MZprox1^-/-^* mutant and *Zprox1^-/-^* mutant datasets filtered aggregated count files were processed, sc-transform normalised, filtered and clustered (louvain) using *Seurat* version 2.0 ^65^ and 3.0 ^34^ respectively for *R statistical software* version 4.0.2. QC was evaluated before and after normalisation using plot functions in *Seurat* and *scater* 1.20.1 ^66^, and all thresholds and settings are described in scripting. Cluster solutions were evaluated using *ClusTree* ^67^. Datasets used in the atlas of lymphangiogenesis were processed, filtered, merged and log-normalised using *Seurat* version 3.0 ^34^, with QC and settings as above. Merged data was clustered and normalised using *CSS simspec* ^33^ and clustering and cluster evaluation performed on this object only, as described above. For all scRNA-seq datasets cluster phenotype was determined using the expression of key markers (Supplementary Table), with the aid of *CellXGene* visualisation software ^68^. All downstream analysis and plotting were performed using *Seurat* version 3.0 ^34^ with default settings. All gene ontology analyses were performed using Panther.db ^37^ (Biological Process Complete).

### Preparation of single nuclei for snATAC-seq

Single cell suspensions were sorted by FACS and prepared for nuclei isolation as previously described by 10x Genomics Demonstrated Protocol for Single Cell ATAC Sequencing (CG000169 - Rev D). Cell suspensions were pelleted (300 x g for 5 minutes) and rinsed with PBS + 0.04% BSA. Cells were resuspended in 95uL of freshly prepared lysis buffer (10mM Tris-HCl pH 7.4, 10 mM NaCl, 3 mM MgCl2, 0.1% Tween-20, 0.1% NP40 Substitute, 0.01% Digitonin, and 1% BSA) and incubated on ice for 1 minute. 100uL of chilled wash buffer (10 mM Tris-HCl, pH 7.4, 10 mM NaCl, 3 mM MgCl2, 0.1% Tween-20, 1% BSA) was used to neutralise the reaction, before the nuclei were pelleted (500 x g for 5 minutes) and resuspended again in 7uL of 1x Nuclei Buffer (10X Genomics Cat# PN-2000153/2000207). Presence of healthy and intact nuclei was assessed by visual inspection on a brightfield microscope using Trypan Blue staining (Thermo Fisher Cat# T10282) and Countess Cell Counting Chamber Slides (Thermo Fisher Cat# C10228).

### snATAC-seq library preparation and sequencing

Single nuclei suspensions were resuspended at approximately 5000 nuclei per μL before undergoing tagmentation for 60min at 37°C. After tagmentation nuclei were partitioned and barcoded using the 10X Genomics Chromium Controller (10X Genomics) and the Single Cell ATAC Reagent Kit (V1.1; 10X Genomics; PN-1000176). Tagmented nuclei were loaded onto the Chromium Single Cell Chip H (10X Genomics; PN-1000162), GEM generation, barcoding and library construction was performed according to the 10X Genomics Chromium User Guide. The resulting single cell ATAC libraries contained unique sample indices for each sample. The libraries were quantified on the Agilent BioAnalyzer 2100 using the High Sensitivity DNA Kit (Agilent, 5067-4626) and pooled in equimolar ratios. Sequencing was performed on an Illumina NextSeq 500 using a 150-cycle High-Output Kit as follows: 50bp (Read1), 8bp (i7 index), 16bp (i5 index), 50bp (Read2) achieving a read depth of 25,000 read pairs per nucleus.

### snATAC-seq processing and analysis

FASTQ files generated from sequencing were used as inputs to 10X Genomics *Cell Ranger ATAC 2.0.0*. *cellranger-atac count* was used to generate count files mapped to GRCz11 (ENSEMBL 101), without depth normalisation. Resulting fragment files were read into *ArchR 1.0.1* for *R statistical software 4.0.5* as a tile matrix with 500-bp bins. All remaining steps in the ATAC-Seq analysis were performed within *ArchR 1.0.1*. QC filtering was performed, and only high-quality cells with a TSS enrichment score greater than 4 and greater than 1,000 unique nuclear fragments were retained. Doublets were predicted using *addDoubletScores* and filtered using *filterDoublets.* Data normalization and dimensionality reduction were performed using iterative Latent Semantic Indexing (LSI) and Uniform Manifold Approximation and Projection (UMAP) embeddings were used for visualisation in reduced dimension space. Separate from the *ArchR 1.0.1* package, cluster solutions were independently evaluated using *clustree 0.4.3*. A Gene Score Matrix that stores predicted gene expression was then generated based on the accessibility of regulatory elements in the vicinity of the gene. We used gene scores of endothelial markers for cluster annotation and subsetting. Differentially accessible genes were identified by differential testing using *getMarkerFeatures*. Local chromatin accessibility of the marker genes was visualised using the *plotBrowserTrack*.

### Gene regulatory network analyses

For gene regulatory network construction we leveraged the ENCODE transcription factor targets gene-attribute edge list from the Harmonize database https://maayanlab.cloud/Harmonizome/dataset/ENCODE+Transcription+Factor+Targets that includes information for n=181 transcription factors from ChIP-seq analyses^42^. Gene lists generated from scRNA-seq or snATAC-seq analysis were mapped from Zebrafish genome version GRCz11 to Human genome version GRCh38.p13 using ENSEMBL Biomart ^69^. These mapped gene sets were used to select relevant edges, that were visualised as a degree-sorted circular network in Cytoscape^70^.

## EXTENDED DATA FIGURES AND FIGURE LEGENDS

**Extended data Fig 1:**
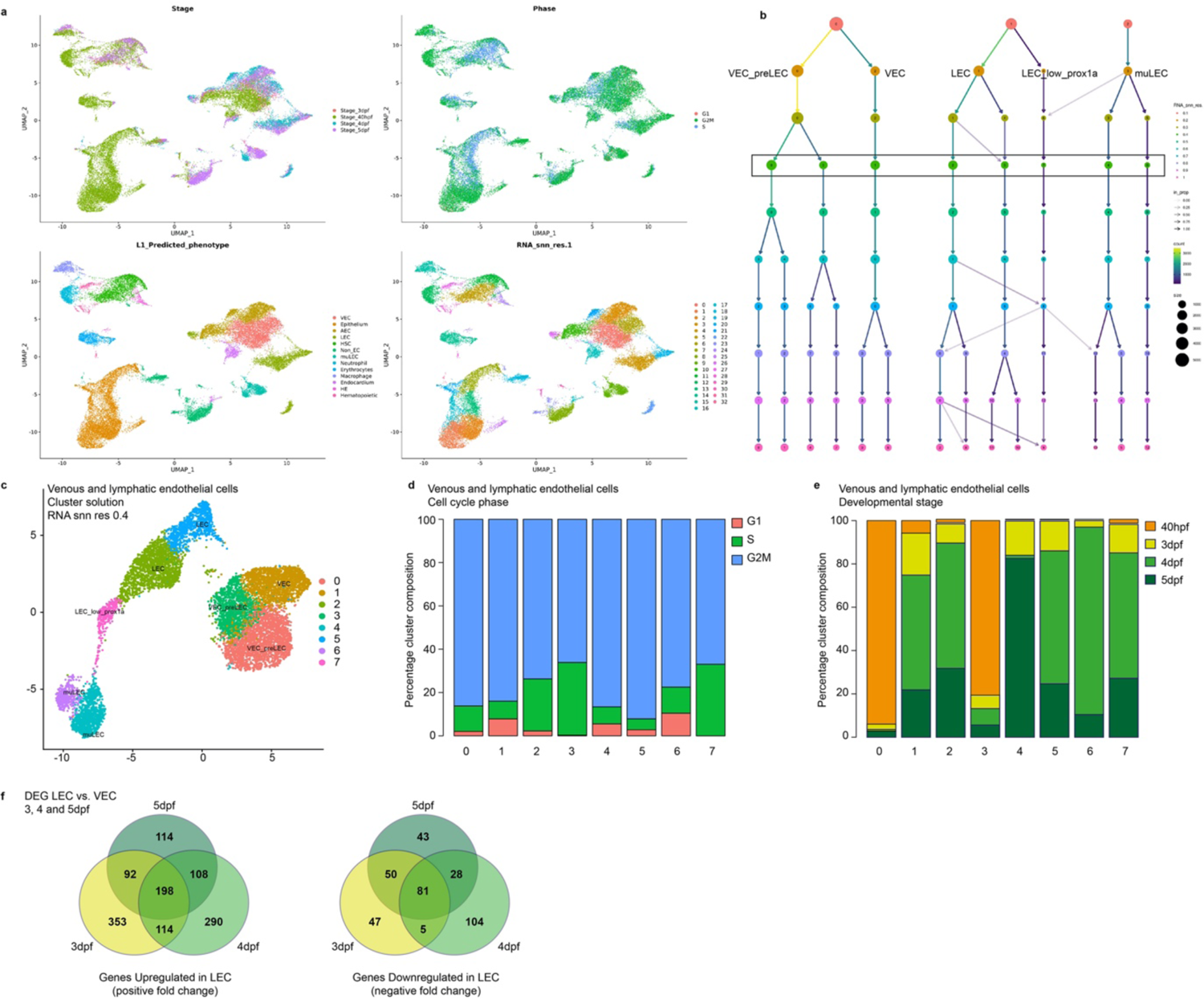
Full dataset and analysis for data in Fig 1. **a.** UMAP visualization of n=35634 endothelial cells sequenced for the single cell atlas of lymphangiogenesis (n=6 samples; see also **Extended data table 1a** for markers used to identify clusters) coloured according to developmental stage (top left), cell cycle phase (top right), predicted cell phenotype (bottom left), and cluster solution (RNA-snn-res.1). **b.** Clustree analysis demonstrating the relationship between different cluster resolutions for the n=9771 venous and lymphatic endothelial cells in Figure 1a. Resolution 0.4 (boxed) was used for all downstream analyses. **c.** UMAP visualization of cluster resolution 0.4 (as appears in Fig 1). **d.** Stacked bar plot of resolution 0.4 cluster composition cell cycle phase. **e.** Stacked bar plot of resolution 0.4 cluster composition developmental stage. **f.** Venn diagram of significant genes commonly upregulated in LEC compared to VEC at 3, 4 and 5dpf.

**Extended data Fig 2:**
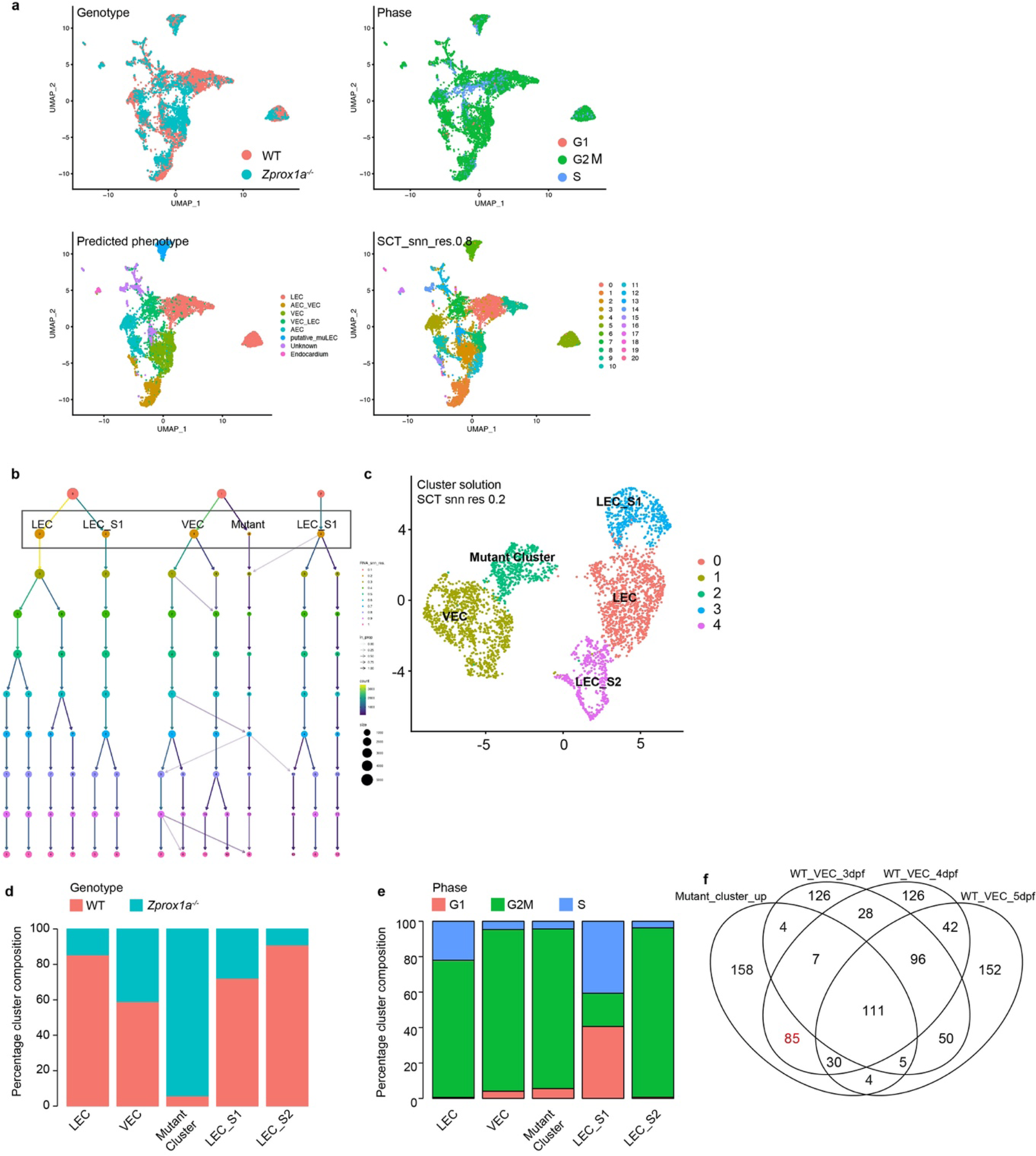
Full dataset and analysis for data in Fig 3. **a.** UMAP visualization of n=8075 endothelial cells sequenced comparing lymphangiogenesis in WT and *Zprox1a* mutants at 4dpf (n=2 samples; see also **Extended data table 2a** for markers used to identify clusters) coloured according to developmental stage (top left), cell cycle phase (top right), predicted cell phenotype (bottom left), and cluster solution (SCT-snn-res.8) **b.** Clustree analysis demonstrating the relationship between different cluster resolutions for the n=3063 venous and lymphatic endothelial cells in Figure 3a. Resolution 0.2 (boxed) was used for all downstream analyses. **c.** UMAP visualization of cluster resolution 0.2 **d.** Stacked bar plot of resolution 0.2 cluster composition genotype **e.** Stacked bar plot of resolution 0.2 cluster composition cell cycle phase **f.** Venn diagram indicates the overlap between genes significantly upregulated in VEC compared to LEC at each developmental stage of the single cell atlas, and those upregulated in the *Zprox1a^-/-^* Mutant Cluster at 4dpf compared to WT LECs. N=85 genes (coloured red) are uniquely associated with the Mutant Cluster and WT VEC at 4dpf, compared to N=4 genes in WT VEC at 3 and 5dpf.

**Extended data Fig 3:**
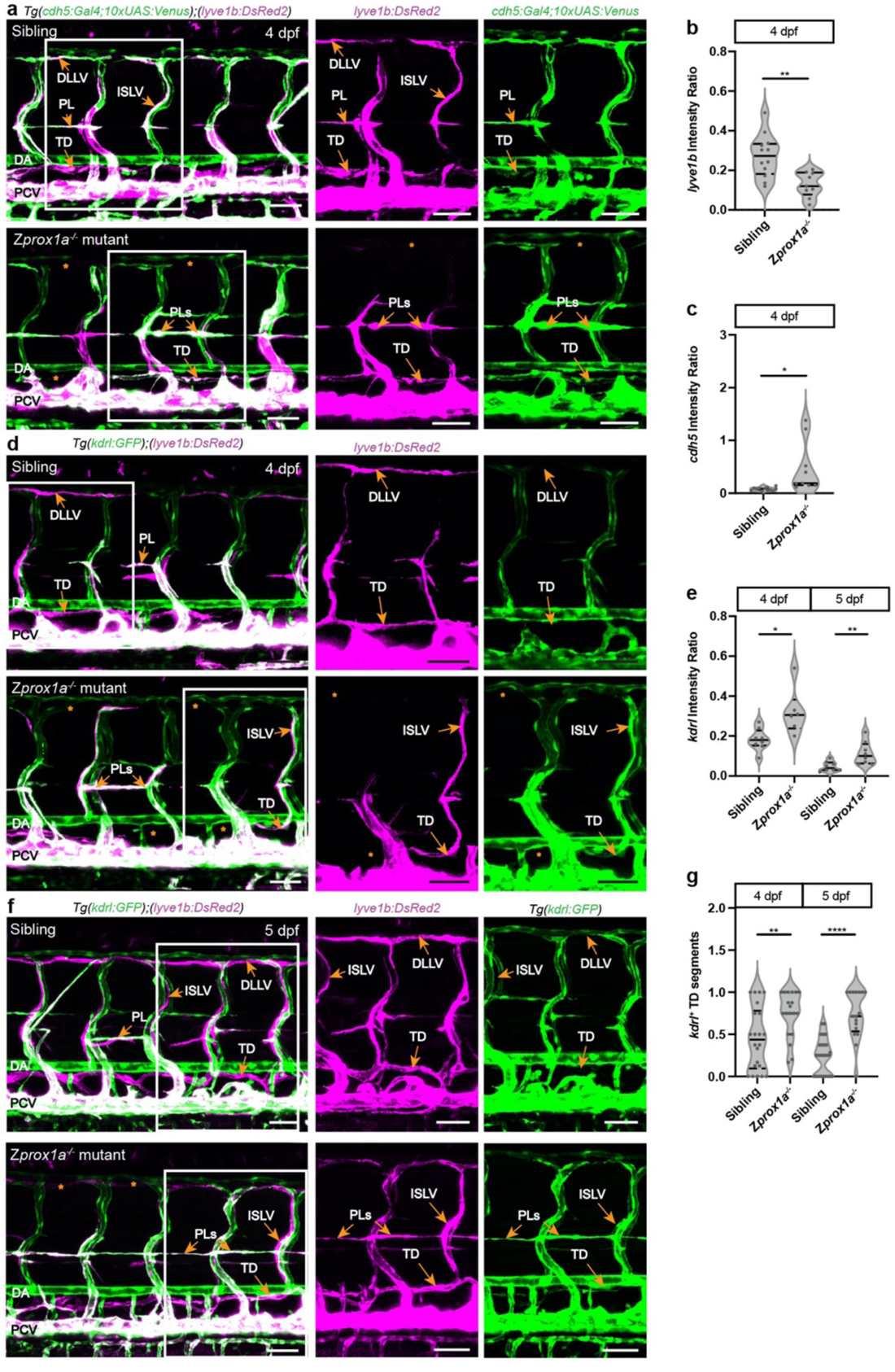
Additional phenotypic analysis of blood vascular markers in *Zprox1a^-/-^* mutants associated with Fig 3 **a.** Lateral confocal images of zebrafish trunks at 4dpf showing *cdh5* expression (green) and *lyve1b* expression in lymphatic vessels (magenta) in control (upper) and *Zprox1a* mutant embryos (lower). Transgenic lines are indicated. Scale bar, 50 µm. **b.** Quantification of *lyve1b* expression intensity in the TD relative to the PCV in sibling and *Zprox1a* mutant embryos at 4dpf. **c.** Quantification of *cdh5* expression intensity in the TD relative to the DA in sibling and *Zprox1a* mutant embryos at 4dpf. **d.** Lateral confocal images of zebrafish trunks at 4dpf showing *kdrl* expression (green) and *lyve1b* expression in lymphatic vessels (magenta) in control (upper) and *Zprox1a* mutants (lower). Transgenic lines are indicated. Scale bars, 50 µm. **e.** Quantification of *kdrl* expression intensity in the TD relative to the DA in sibling and *Zprox1a* mutant embryos at 4dpf and 5dpf (upper). **f.** Lateral confocal images of zebrafish trunks at 5dpf showing *kdrl* expression (green) and *lyve1b* expression in lymphatic vessels (magenta) in control (upper) and *Zprox1a* mutants (lower). Transgenic lines are indicated. Scale bars, 50 µm. **g.** Quantification of the number of TD segments displaying *kdrl* expression in *Zprox1a* mutant embryos at 4dpf and 5dpf (lower). DLLV, dorsal longitudinal lymphatic vessel. ISLV, intersomitic lymphatic vessel. TD, thoracic duct. DA, Dorsal aorta. PCV, posterior cardinal vein. vISV, venous intersegmental vessel. PL, Parachoral lymphatic endothelial cells.

**Extended data Fig 4:**
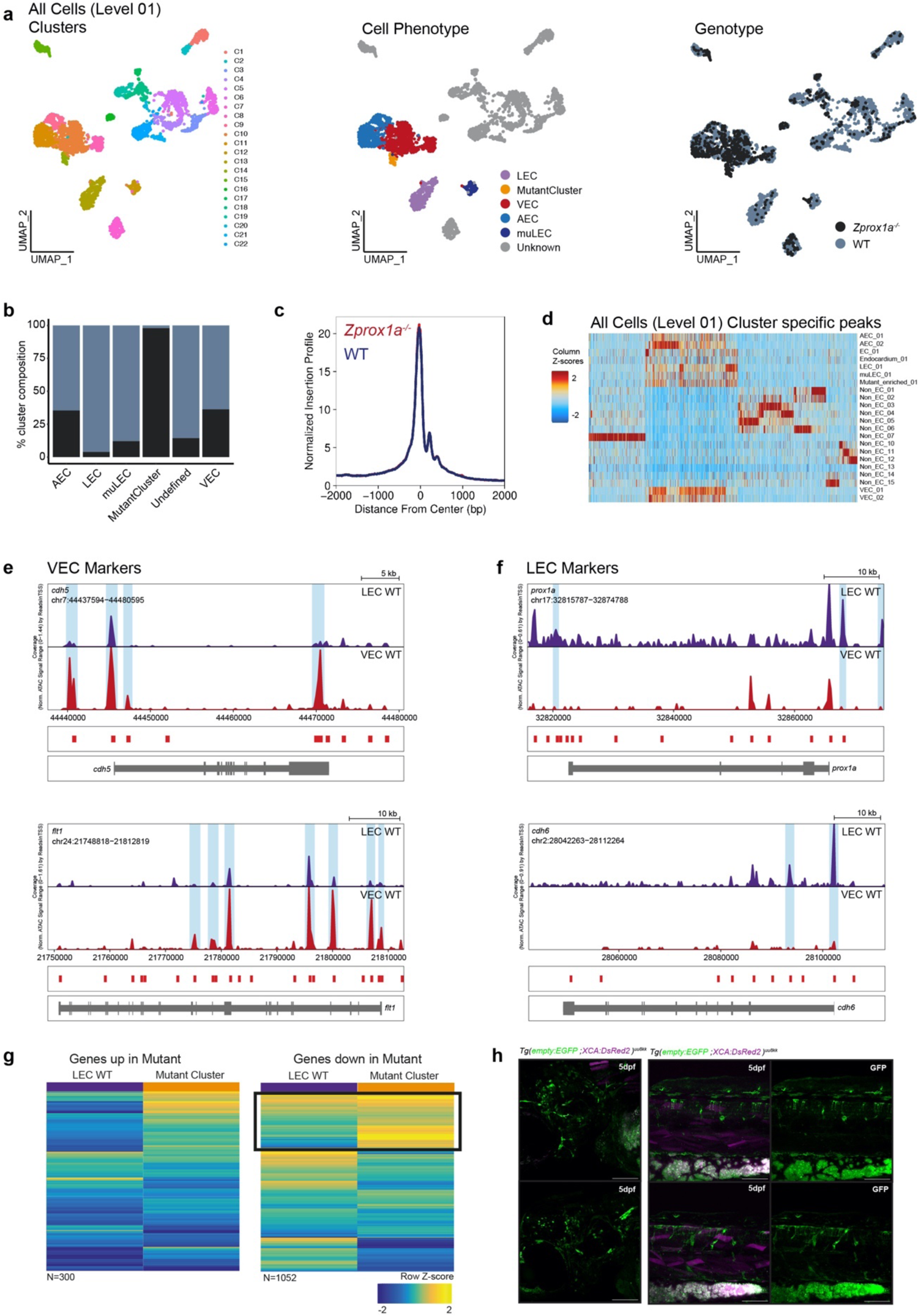
Additional data and analysis for data in Fig 4. **a.** UMAP visualization of n=3,731 single nuclei ATAC-seq data comparing lymphangiogenesis in WT and *Zprox1a* mutants at 4dpf. Nuclei coloured according to cluster (Res-0.8, left), predicted cell phenotype (middle) and genotype (right). Cell phenotype was predicted based on accessibility changes at the genes defined in **Extended data table 3a. b.** Stacked bar plot indicating the genotype composition of nuclei captured per cell type. **c.** Density plot indicating the majority of snATAC-seq peaks fall around the TSS. Y-axis represents distance from TSS (bp) and y-axis represents normalised Tn5 insertion profile. **d.** Heatmap of cluster specific accessible peaks for all cell clusters. Colour reflects a column-wise Z-score, each row represents a cluster defined in **Extended Data Fig. 4a**, columns are peaks. **e.** Genome accessibility track of key VEC markers *cdh5* (top) and *flt1* (bottom) indicating these genes are more accessible in WT VEC than WT LEC cell phenotypes. Red bars represent peaks in the reproducible peak set from snATAC-seq. DAP between WT VECs and WT LECs (Wilcoxon Rank Sum, *FDR* < 0.05) are highlighted blue. **f.** Genome accessibility track of key LEC markers *prox1a* (top) and *cdh6* (bottom) indicating these genes are more accessible in WT LEC than WT VEC cell phenotypes. Red bars represent peaks in the reproducible peak set from snATAC-seq. DAP between WT LECs and WT VECs (Wilcoxon Rank Sum, *FDR* < 0.05) are highlighted blue. **g.** Heatmaps of accessibility (gene score) for DEG (Fig 3f) upregulated (left, n=300 mapped genes) and downregulated (right, n=1,052 mapped genes) in the Mutant Cluster compared to WT LECs. Colour indicates row-wise Z-score, each row represents a differentially expressed gene in scRNA-seq data, columns are snATAC-seq clusters. Boxed genes downregulated in Mutant display more permissive chromatin in Mutant than WT LEC, showing that mutant cluster cells display a unique chromatin state at many genes. **h.** Confocal projections of zebrafish head (left) labelled with *Tg(empty:EGFP; XCA:DsRed2)* at 5 dpf, showing GFP expression in neuronal tissues. Confocal projections of zebrafish trunk (right) labelled with *Tg(empty:EGFP; XCA:DsRed2)* at 5 dpf, showing GFP expression in neuronal tissues. All scale bars = 100μm.

**Extended data Fig 5:**
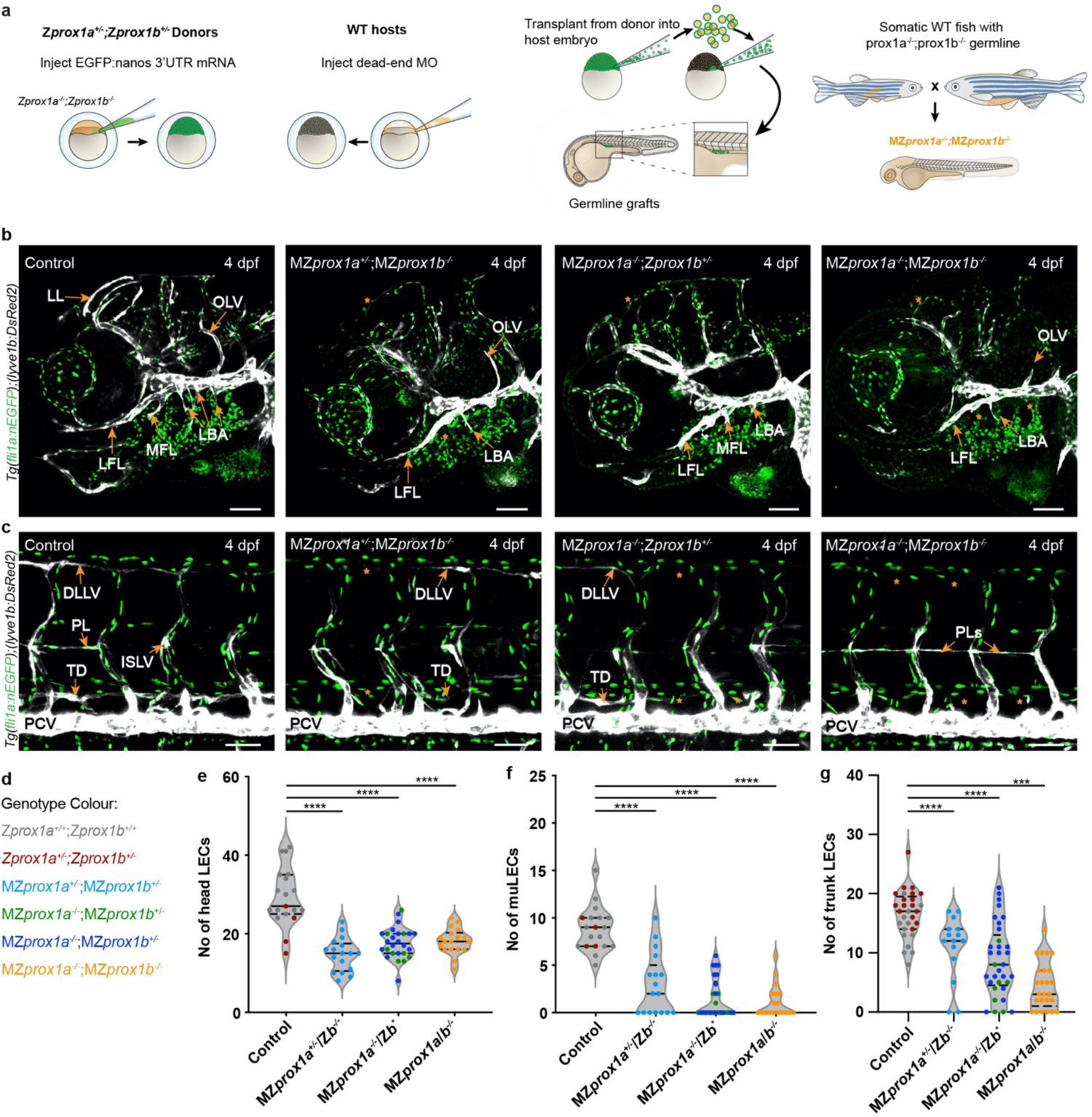
Additional phenotypic analysis of *MZprox1a^-/-^;MZprox1b^-/-^* mutants associated with Fig 6 **a.** Schematic explanation of the germline replacement transplantation method used to generate *MZprox1a^-/-^;MZprox1b^-/-^* mutants. **b-c.** Lateral confocal images showing vascular defects in control and mutant in the craniofacial (**b**) and trunk (**c**) regions of the larvae at 4dpf. Genotypes and transgenic labels are indicated. *MZprox1a-/-* and *MZprox1a+/-, MZprox1b-/-* animals both showed a strong loss of lymphatic vessels in craniofacial and trunk regions. The *MZprox1a-/-, MZprox1b-/-* mutants showed a robust loss of craniofacial vessels equivalent to the other mutant phenotypes but a more severe loss of trunk lymphatics and an accumulation of PLs in the HM. **d.** Genotypes that are shown in the quantification. Colour codes are used to identify individual embryonic genotypes in the dot plots showing quantification in **e-g**. **e.** Quantification of the number of craniofacial (head) LECs within facial lymphatic vessels showing evidence for contributions from *prox1a* and *prox1b*. **f.** Quantification of the number of mural LECs (muLECs) showing a major role for *prox1a* and a role for *prox1b* in muLEC development. **g.** Quantification of the number of trunk LECs within vessels showing a dominant role for *prox1a* in trunk lymphangiogenesis. Z, zygotic; MZ, maternal and zygotic; HM, horizontal myoseptum; LL, lymphatic loop; OLV, otholitic lymphatic vessel; LFL, lateral facial lymphatic vessel; MFL, medial facial lymphatic vessel; LBA, lymphatic branchial arches; DLLV, dorsal longitudinal lymphatic vessel; PL, parachordal LEC; ISLV, intersomitic lymphatic vessel; TD, thoracic duct; DA, dorsal aorta; PCV, posterior cardinal vein.

**Extended data Fig 6:**
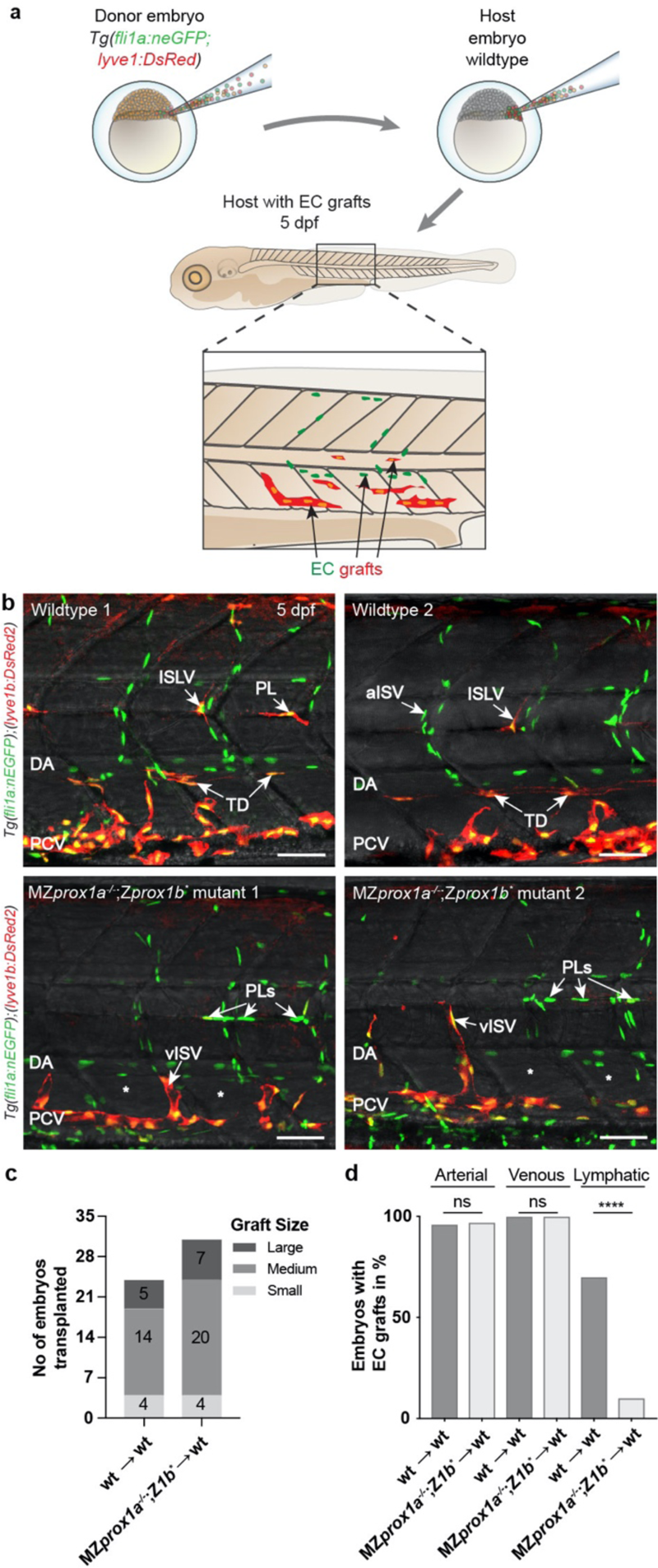
Embryonic transplantation data demonstrating endothelial cell autonomy of *prox1a* maternal zygotic mutant phenotype associated with Fig 6. **a.** Schematic showing the endothelial cell transplantation approach used to test cell autonomy for the vascular phenotypes seen in *MZprox1a* mutants. Cell identities were traced following transplantation in unlabelled host embryos using expression of the pan-endothelial marker *fli1a* and venous/lymphatic marker *lyve1b* (indicated). **b.** Examples of 5dpf wildtype hosts engrafted with wildtype donor cells (upper) that contribute to arteries (aISVs, DA), veins (PCV, vISVs) and lymphatic vessels (TD, ISLVs) and examples of wildtype embryos engrafted with *MZprox1a* mutant cells contributing arteries (aISVs, DA) and veins (vISV, PCV) but not lymphatic vessels. Note the PL accumulation in the horizontal myoseptum.. **c.** The number of embryos imaged and graft sizes scored post-tranplantation. As the contribution to the different EC types is affected by the graft size, we analysed grafts of comparable sizes and position from wildtype and mutant embryos. Grafts were categorised as large, medium or small. Small (spanning 1-2 somites), medium (spanning 2-3 somites) and large grafts (spanning 4-5 somites). The number of ECs contributing to each EC type was scored at single cell resolution and the percentage of grafts with a contribution to each lineage calculated accordingly. The numbers within the different graft groups indicate the number of analysed embryos within that group. **d.** The percentage of embryos with vascular grafts that display contributions to arteries, veins and lymphatics. *MZprox1a^-/-^* mutant cells contribute to arteries and veins but showed a near complete loss of contribution to lymphatics. The size of the grafts did not influence the outcome. *Z1b\** indicates a mixed prox1b background. p < 0.0001 from an unpaired, two-sided t-test. DA, dorsal aorta. PCV, posterior cardinal vein. PL, parachordal LEC. TD, thoracic duct. aISV, arterial intersegmental vessel. vISV, venous intersegmental vessel. ISLV, intersegmental lymphatic vessel.

**Extended data Fig 7:**
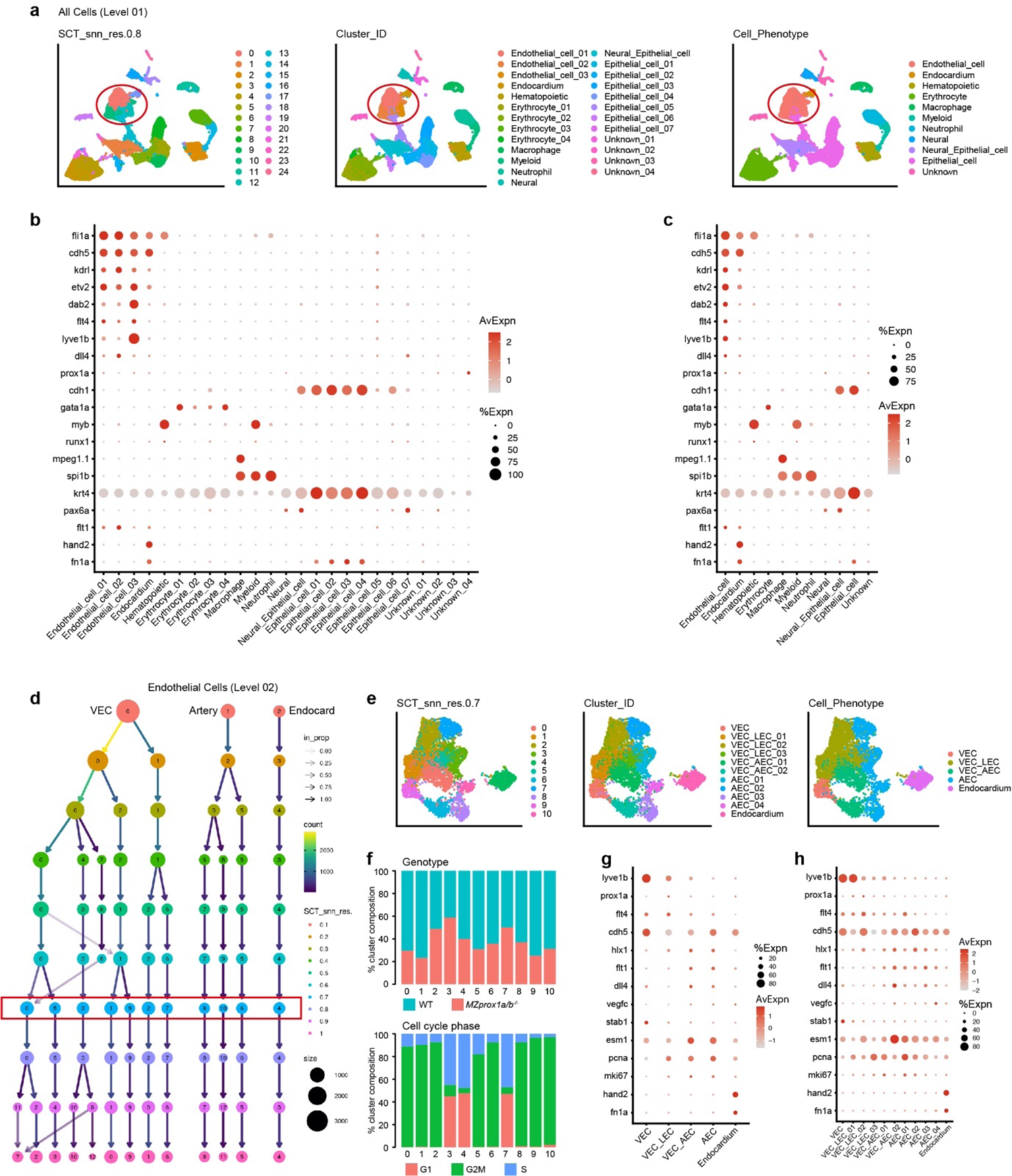
Additional data and analysis for data in Fig 7. **a.** UMAP visualization of n=26,289 endothelial cells sequenced comparing lymphangiogenesis in WT and *MZprox1a/b* mutants at 40hpf (n=4 samples; see also **Extended data table 5a** for markers used to identify clusters) coloured according to cluster solution (SCT-snn-res.8, left), predicted phenotype per cluster (middle), and predicted cell phenotype (right). Endothelial cells selected for further analysis are indicated by a red circle. **b.** Dot plot summarizing the scRNA-seq expression level of n=20 key genes used to predict cell phenotypes of each cluster in the complete dataset (Level 01), for each cluster defined in **Extended Data Fig. 7a** (SCT-snn-res.8). The size of the dot represents the proportion of cells that express the markers in the cluster, and colour represents SCT-normalised expression. **c.** Dot plot summarizing the scRNA-seq expression level of n=20 key genes used to predict cell phenotype in the complete dataset. The size of the dot represents the proportion of cells that express the markers in the cluster, and colour represents SCT-normalised expression. **d.** Clustree analysis demonstrating the relationship between different cluster resolutions for the n=5,347 endocardial, venous and arterial endothelial cells in Fig. 7a and **Extended Data Fig. 7a** (Level 02). Cluster resolution SCT_snn_0.7 (boxed) was selected for all downstream analyses. **e.** UMAP visualization of cluster resolution SCT_snn_0.7 (left), predicted phenotype of each cluster (middle) and phenotypic group (right). **f.** Stacked bar plot of cluster resolution SCT_snn_0.7 cluster composition genotype (top) and cell cycle phase (bottom). **g.** Dot plot summarizing the scRNA-seq expression level of n=14 key genes used to predict cell phenotype in the endothelial cell populations (Level 02, n=5,347 cells). The size of the dot represents the proportion of cells that express the markers in the cluster, and colour represents SCT-normalised expression.

